# Evolutionarily recent transcription factors partake in human cell cycle regulation

**DOI:** 10.1101/2024.11.04.621792

**Authors:** Cyril Pulver, Romain Forey, Alex R. Lederer, Martina Begnis, Olga Rosspopoff, Joana Carlevaro-Fita, Filipe Martins, Evarist Planet, Julien Duc, Charlène Raclot, Sandra Offner, Alexandre Coudray, Arianna Dorschel, Didier Trono

## Abstract

The cell cycle is a fundamental process in eukaryotic biology and is accordingly controlled by a highly conserved core signaling cascade. However, whether recently evolved proteins also influence this process is unclear. Here, we systematically map the influence of evolutionarily recent transcription factors (TFs) on human cell cycle progression. We find that the genomic targets of select young TFs, many of which belong to the rapidly evolving Krüppel-associated box (KRAB) zinc-finger proteins (KZFP) family, exhibit synchronized cell cycle expression. Systematic perturbation studies reveal that silencing recent TFs disrupts normal cell cycle progression, which we experimentally confirm for ZNF519, a simian-restricted KZFP. Further, we show that the therian-specific KZFP ZNF274 sets the cell cycle expression and replication timing of hundreds of clustered genes. These findings highlight an underappreciated level of lineage specificity in cell cycle regulation.

## Introduction

The cell cycle is the universal process allowing eukaryotic cells to reproduce. Its completion reflects progression through a highly conserved core signalling cascade, as evidenced by complementation experiments performed across deeply diverged organisms (Draetta *et al.*, 1987; Lee and Nurse, 1987; Jimenez *et al.*, 1990). Whether it warrants generalizing conservation to the cell cycle as a whole has however been subject to debate ^1–3^. For instance, metazoans and fungi achieve cell cycle entry through analogous positive feedback loops carried out by seemingly unrelated proteins. Thus, molecular network topology - rather than the molecular players themselves - has been put forth as evolutionarily conserved ^2,3^. This view leaves room for substantial levels of lineage specificity whereby recently emerged proteins may partake in regulating the cell cycle while leaving the overarching logic unaffected ^2–4^. However, the extent of that phenomenon remains poorly understood. This partially stems from the broad use of conservation as a proxy for cell cycle function which, while pertinent to single out candidates for functional studies, leaves a blind spot on evolutionarily recent proteins.

Here, we combine forward genetics with genomics, mathematical modelling and functional assays to study the role of evolutionarily recent transcription factors (TFs) in human cell cycle progression. Specifically, we establish a synchronization-free chronogram of cell cycle expression to reveal that genomic targets of evolutionarily recent TFs – many belonging to the fast-evolving family of Krüppel-associated box (KRAB) zinc-finger proteins (KZFPs) – display synchronized cell cycle expression. We then systematically map the impact of gene silencing on cell cycle progression by re-analyzing a large-scale perturbation dataset whereby all genes expressed in human leukemia K562 cells were individually silenced, observing that some evolutionarily recent TFs – including KZFPs - are required for unaltered cell cycle progression. We confirm this experimentally for ZNF519, a KZFP restricted to simians. Lastly, we show that the mammalian-specific KZFP ZNF274 sets the replication timing and cell cycle expression of hundreds of clustered genes, including most *KZFPs* themselves. These functional dependences having emerged in the last 20 to 150 million years of mammalian evolution, our work suggests previously unappreciated levels of lineage specificity in cell cycle regulation.

## Results

### Evolutionarily recent TFs display binding profiles suggestive of cell cycle regulation

We sought to gauge the cell cycle regulatory potential of evolutionarily recent human TFs by systematically assessing their binding enrichment at the promoters of cell cycle-rhythmic genes. To this end, we first established a synchronization-free cell cycle chronogram of gene expression. More specifically, we leveraged an RNA-preserving fixation technique ^5^ to FACS-sort, based on CCNA vs. DAPI, untreated K562 into eight cell cycle fractions, which we interrogated by RNA-seq (Fig. 1A, S1A). We tested 13,749 expressed protein-coding genes for cell cycle rhythmicity by modeling their expression as sinusoids ^6^ (e.g. : Fig. S1B), thus identifying 4,894 rhythmic genes (adj. p < 0.05, F test) along with their baseline expression and phase of peak expression (acrophase) (Fig. 1B), available for exploration at https://tronoapps.epfl.ch/cycleQuest/. We compared acrophases against cell cycle gene harmonics computed on scRNA-seq data emanating from untreated K562, HL60, SUDHL4 and THP1 cells ^7^ using *VeloCycle* ^8^, finding a remarkable consistency (Fig. S1C-D). Moreover, rhythmic genes were enriched for phase-specific cell cycle processes, e.g. DNA replication initiation in G1/S-to-S1 followed by establishment of chromosome localization in G2 and mitotic cytokinesis in G2/M (Fig. S1E); and many of the top rhythmic genes were known cell cycle effectors, such as *MCM4/6*, *CHAF1B* and *DONSON* (Fig. 1B). In general, genes peaking in M-to-S1 – thereafter referred to as G1-centered (G1-c) genes - were expressed at lower baseline levels and replicated later than their S2-to-G2/M – G2- centered (G2-c) - counterparts (Fig. 1B-D, Fig. S1F,G), accordingly displaying stronger binding by the late replication timing factor RIF1 ^9,10^ (Fig. 1B, E, Fig. S1H) and more elevated H3K9me3 promoter levels (Fig. 1B, F, Fig. S1I). We thus recovered canonical features of the cell cycle transcriptional program, namely correlations between expression, rhythmicity, replication timing and chromatin states. Together, these results justified leveraging our bulk RNA-seq chronogram, made available for further exploration (https://tronoapps.epfl.ch/cycleQuest/), for the characterization of cell cycle regulation.

**Figure 1:**
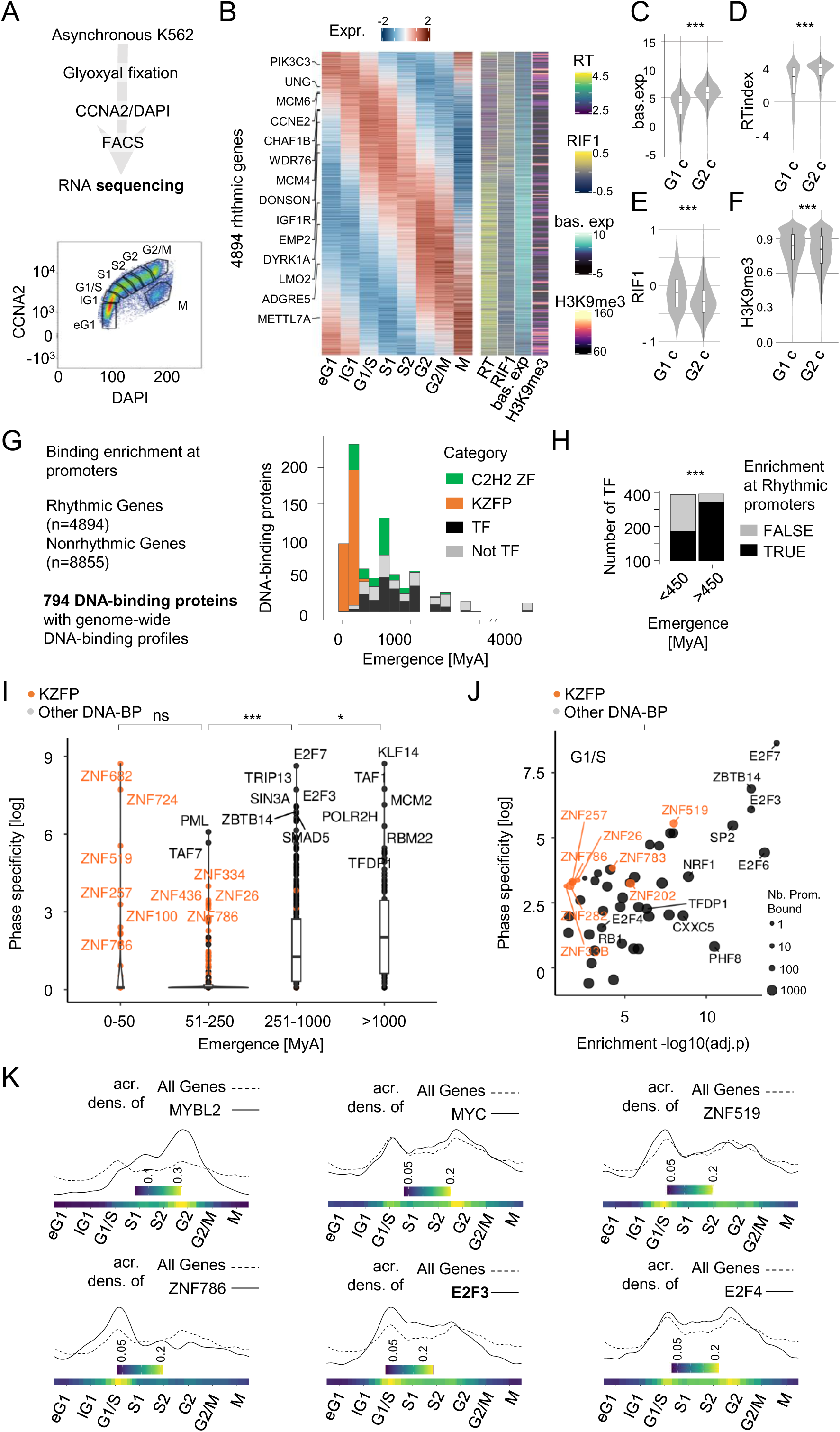
Evolutionarily recent TFs display binding profiles suggestive of cell cycle regulation. (A) Top: overview of the synchronization-free procedure for establishing the cell cycle chronogram of gene expression. K562 cells were fixed (Channathodiyil and Houseley, 2021), FACS-sorted, followed by RNA extraction and sequencing. Bottom: FACS-sorted K562 stained with DAPI (x-axis) and CCNA2 antibody (y-axis). Sorting gates are annotated in black. (B) Left: z-scored expression of rhythmic (adj. p-val. < 0.05, F-test) genes along the eight cell cycle bins defined in Fig. 1A. Rhythmicity is tested by fitting the expression of each gene to a sinusoid (Cornelissen, 2014). The top two rhythmic genes (smallest adj. p-val.) peaking in each bin are highlighted. Right: 1) Replication Timing (RT) index of K562 (Hansen *et al.*, 2010), 2) log2 RIF1 binding enrichment over input (Klein *et al.*, 2021), 3) estimated baseline gene expression, and 4) H3K9me3 promoter coverage (The ENCODE Project Consortium, 2012). (C) Baseline expression, (D) RT index, (E) RIF1 binding and (F) H3K9me3 promoter coverage of rhythmic genes peaking in G1-centered phases (M to S1, G1-c) or G2-centered phases (S2 to G2/M, G2-c). (G) Left: Enrichment of DNA-binding proteins (DBP) at promoters of rhythmic genes. Promoters of rhythmic genes peaking in each phase (eG1, lG1, G1/S, S1, S2, G2, G2/M, M) were checked for enrichment (adj. p-val. < 0.05, Fisher’s Exact Test) of DBPs (The ENCODE Project Consortium, 2012; Najafabadi *et al.*, 2015; Schmitges *et al.*, 2016; Imbeault, Helleboid and Trono, 2017; Helleboid *et al.*, 2019; The ENCODE Project Consortium *et al.*, 2020; De Tribolet-Hardy, 2022). Right: Emergence (Imbeault, Helleboid and Trono, 2017; Shao *et al.*, 2019; Tong *et al.*, 2021; De Tribolet-Hardy, 2022) of human DBPs throughout evolution. (H) DBPs enriched (adj. p < 0.05, Fisher’s Exact Test) at rhythmic promoters in at least one phase, split according to their evolutionary age. Enrichment of old (>450myo) DBPs amongst rhythmic promoters-enriched DBPs was computed using Fisher’s Exact Test. “***” indicates p < 0.005. (I) Phase specificity of DBP enrichment at rhythmic promoters (i.e. median rank of Fisher’s Exact Test adj. p-value across the eight phases divided by the rank of adj. p-value in the phase of interest) relative to their evolutionary emergence. The most specific DBPs in each age category are highlighted, with KZFPs in orange. (J) DBP enrichment (-log10(adj. p-value)) and specificity across rhythmic promoters reaching peak expression in G1/S. Dot area is proportional to the number of genes whose promoter is bound the DBP. KZFPs are highlighted in orange.

We next wondered whether evolutionarily recent TFs were enriched at the promoters of phase-specific rhythmic genes. To this end, we aggregated genome-wide DNA binding profiles assayed in K562 by ENCODE ^11,12^ with those obtained in HEK293T for the rapidly expanding family of C2H2 zinc finger proteins ^13–17^, which encompasses most human TFs having emerged in the mammalian lineage ^18^ (Fig. 1G). Promoters of synchronous rhythmic genes were enriched for DNA-binding proteins involved in phase-specific processes (Fig. S1J, K), e.g. E2Fs and TFDP1 in G1/S and S phase-specific processes. Thus, this methodology aptly assigns well-known cell cycle effectors to their phase of predominant activity.

We next inspected the age distribution of DNA-binding proteins ^15,17,19,20^ with respect to their enrichment at the promoters of rhythmic genes. Older DNA-binding proteins were overall more frequently enriched at rhythmic genes (Fig. 1H) with greater phase specificity, i.e. high enrichment at the promoters of rhythmic genes peaking in one phase relative to those of rhythmic genes peaking in other phases (Fig. 1I). These observations illustrate the prevalence of deep conservation in cell cycle progression. We nonetheless found intriguing occurrences of evolutionarily recent TFs displaying substantial and phase-specific enrichments at the promoters of rhythmic genes (Fig. 1I). Among these, many belonged to the fast-evolving family of Krüppel-associated box (KRAB) zinc-finger proteins (KZFPs) ^15–17,21^. For instance, the ∼43-million-year-old (myo) and primate-specific ZNF682 and ZNF724 were specifically enriched at promoters of lG1-peaking genes (Fig. S1K). Moreover, promoters of G1/S rhythmic genes were highly enriched for ZNF519 (Fig. 1J, S1K), a 20myo Hominidae-specific KZFP recently linked to the evolution of promoter sequences ^22^. ZNF519 scored comparably to the deeply conserved cell cycle master regulators E2F3 and E2F6 in G1/S, both in terms of rhythmic promoter enrichment and phase specificity, even superseding RB1, E2F4 and TFDP1 (Fig. 1J). Together, these data suggest that recently emerged TFs, specifically KZFPs, may transcriptionally regulate cell cycle rhythmic genes.

### Perturbing evolutionarily recent TFs causes cell cycle imbalances

To tackle these points, we first assessed the impact of silencing individual genes on the cell cycle distribution of K562 cells. To this end, we leveraged *VeloCycle* to compute cell cycle phases for 1.97 million cells drawn from a perturb-seq experiment whereby 9,326 expressed genes were individually silenced through CRISPRi ^23^. We compared perturbation-induced cell cycle distributions against a control distribution of 75,328 cells transduced with non-targeting gRNAs (Fig. S2A) and projected the resulting relative cell cycle phase densities on a UMAP (Fig. 2A, S2B). We found 4,026/9,326 (43%) CRISPRi conditions inducing cell cycle phase-specific imbalances, i.e. cell accumulations or attritions relative to control cells (p < 0.05, see Methods), made available for exploration at https://tronoapps.epfl.ch/cycleQuest/. Overall, genes involved in DNA replication and chromosome organization were enriched amongst those leading to phase imbalances (Fig. S2C). Perturbations leading to extreme imbalances (n = 129, p <= 1e-5) targeted core effectors of cell cycle progression (cyclins, *TFDP1*), DNA replication (MCM subunits, *RFC4*, *GINS4*, *WDHD1*), DNA damage repair (*ATRIP*, *RAD21, RINT1, RNF138, TTI1*) and/or mitosis (*MAD2L1*, *SMC1A*); all of which emerged over 250 million years ago (mya) (Fig. 2B, S2B, S2D). Extreme imbalances otherwise encompassed perturbations against genes essential for RNA transcription/processing (mediator subunits), mitochondrial function (*COX15*, *MRPL19*), translation (*QARS*) and cofactor synthesis (*CIAPIN1, NUBP1*) (Fig. S2B). We noted that perturbations targeting components of the same complex (*GINS1- 4*, *MCM2-3*) or genes involved in the same signaling cascade (*ATR*/*ATRIP*) often co-localized on the UMAP (Fig. S2E), consistent with the induction of similar cell cycle phenotypes (Fig. 2A). Indeed, we observed phase-specific enrichments in imbalance-inducing CRISPRi targets; e.g. gRNAs targeting known DNA replication components were over-represented amongst those leading to attritions in lG1- to-G1/S and accumulations in G2/M (Fig. S2F-G). Together, these results suggest that CRISPRi-induced imbalances either stem from perturbations affecting the cell cycle or relatively orthogonal yet essential processes required for overall fitness. Thus, this rich data, made available for further inquiry (see Methods), allows for the detection and mapping of factors important for cell-cycle regulation, with the caveat that knocking down genes involved in other essential processes may produce resembling imbalance phenotypes.

**Figure 2:**
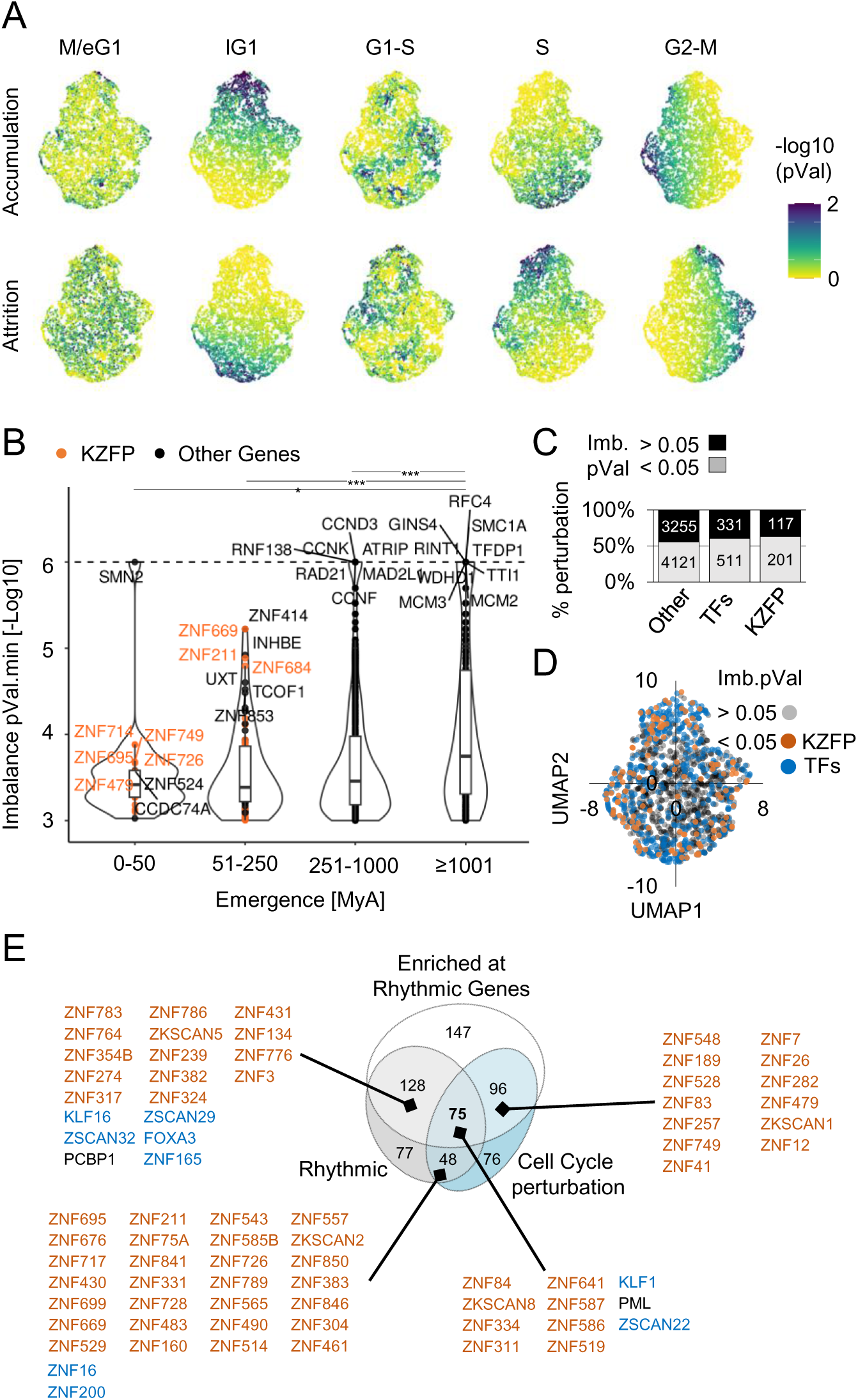
Perturbing evolutionarily recent TFs causes cell cycle imbalances. (A) UMAP projection of cell cycle imbalances computed from a K562 perturb-seq (Replogle *et al.*, 2022) using *VeloCycle* (Lederer *et al.*, 2024). Each dot corresponds to one perturbation (i.e. the silencing of one gene) and is coloured by the statistical significance of accumulation/attrition tests based on a reference multinomial with distribution parameters derived from a binned, unperturbed K562 cell cycle phase distribution (Fig. S2A). (B) Statistical significance of the most severe imbalance (p < 0.05) induced by each perturbation, grouped by evolutionary age. Genes inducing the most profound (smallest p-value) imbalances in each age group are highlighted in orange (*KZFPs*) or black (other genes). (C) Percentage of silenced *KZFPs*, other transcription factors (TFs) and other genes inducing statistically significant imbalances. (D) Same UMAP as in (A), highlighting perturbations targeting *KZFPs* or other TFs inducing statistically significant imbalances. (E) Overlap between: 1/ DBPs causing cell cycle imbalances (p < 0.05) 2/ DBPs enriched at the promoters of rhythmic genes 3/ DBPs encoded by significantly rhythmic genes. KZFPs found in at least 2 categories are annotated in orange. Other young DBPs (<250myo) found in at least 2 categories are annotated in blue.

Interestingly, *ZNF714*, *ZNF695*, *ZNF649*, *ZNF211*, *ZNF684* and *ZNF33A* – all *KZFPs* - were amongst the most evolutionarily recent (0-50myo) yet cell cycle-disruptive genes, which was also true for *ZNF669*, *ZNF211* and *ZNF684* amongst 51-250myo genes (Fig. 2B). Overall, we identified 117/324 (36%) *KZFPs* leading to imbalances when perturbed, a fraction comparable to that of other TFs (331/842, 39%) and other genes (3255/7376, 44%) (Fig. 2C). Moreover, *KZFP* silencing induced diverse imbalance phenotypes, suggestive of individually distinct effects on cell cycle progression (Fig 2D). To pinpoint putative cell cycle-involved TFs, we refined the list of imbalance-inducing DNA-binding proteins using two additional criteria: enrichment at the promoters of cell cycle rhythmic genes and rhythmic cell cycle expression (Fig. 2E). DNA binding proteins satisfying these three criteria were enriched for both cell cycle (e.g. E2F8, TFDP1, RAD51) and immune development (e.g. RUNX1, KLFs) genes (Fig. S2H), suggesting that our systematic approach to uncovering cell cycle-partaking TFs sheds light on functionally relevant molecular players. We found eleven candidate cell cycle- influencing DNA binding proteins, the genes of which emerged in the last 250my. Aside from the cell cycle-associated TRIM-family protein PML, all were C2H2-containing zinc finger proteins (C2H2- ZFPs) (Fig. 2E). One of them, the well-known erythroid development factor KLF1, is likely more ancient, as a fly ortholog with 78% amino acid identity exists ^24^. Eight were KZFPs (ZNF311, ZKSCAN8, ZNF334, ZNF514, ZNF84, ZNF519, ZNF586 and ZNF587) the last three primate-specific. All eight KZFPs bore standard KRAB domains suggestive of a repressor function except ZKSCAN8 ^25^, which does not interact with the canonical KRAB co-factor TRIM28 ^16^ and additionally carries a SCAN domain. We also identified the SCAN C2H2-ZFP ZSCAN22, which is phylogenetically related to KZFPs ^15^. Overall, our integrative analysis of rhythmic cell cycle expression, DNA-binding proteins at rhythmic promoters and CRISPRi-induced imbalances suggests that evolutionarily recent TFs - in particular KZFPs - partake in human cell cycle regulation.

### *ZNF519* exerts a primate-specific layer of cell cycle regulation

We next set out to experimentally assess the impact of evolutionarily recent TFs on cell cycle progression. Unlike other KZFPs found to be highly enriched at rhythmic promoters (e.g. ZNF202, ZNF282), ZNF519 expression itself was rhythmic (Fig. 3A) and its ablation via CRISPRi was associated with a G2/M attrition in the perturb-seq data (Fig. 3B). We confirmed this by depleting ZNF519 in K562 - this time via shRNA - which indeed resulted in decreasing the fraction of cells in G2 (Fig. S3A, Fig 3C). We then ablated ZNF519 rhythmicity via dox-induced overexpression (Fig. 3D). This significantly reduced the proportion of replicating cells as well as levels of EdU incorporation in S (Fig 3E), correlating with decreased proliferation rates relative to the dox-induced LACZ control (Fig 3F). We next interrogated ZNF519-overexpressing and ZNF519-depleted K562 cells by RNA-seq, observing a global downregulation and upregulation, respectively, of genes marked by ZNF519 binding, consistent with a repressor effect (Fig. 3G, H). Genes that were downregulated upon overexpression and upregulated upon knockdown were enriched for ZNF519 binding at their TSS (Fig. 3I). We thus defined high-confidence (HC) ZNF519 targets as genes marked by ZNF519 binding at their TSS, downregulated upon ZNF519 overexpression and upregulated upon ZNF519 knockdown. We identified 501 such targets (Fig. 3I), including 87 that were both rhythmic and required for unimpeded cell cycle progression (Fig. 3J). Among these, we found several key regulators of the cell cycle, including *CDC25A*, *CDC25B*, *MYB*, *GINS2*, *DONSON*, and *CEBPG* (Fig 3K). Interestingly, we found that E2F1, TFDP1 and RB1 colocalized at the *ZNF519* promoter, likely contributing to its rhythmic expression (Fig S3B, C). Lastly, *ZNF519* ranked fourth behind *ZNF695*, *ZNF724P* and *ZNF93* amongst *KZFPs* whose expression correlated positively with proliferation across 33 TCGA cancer types (Fig. S3D), pointing towards a general association between proliferation, i.e. repeated cell cycling, and elevated ZNF519 expression. Together, these results suggest that despite its recent emergence, ZNF519 transcriptionally regulates key cell cycle genes.

**Figure 3:**
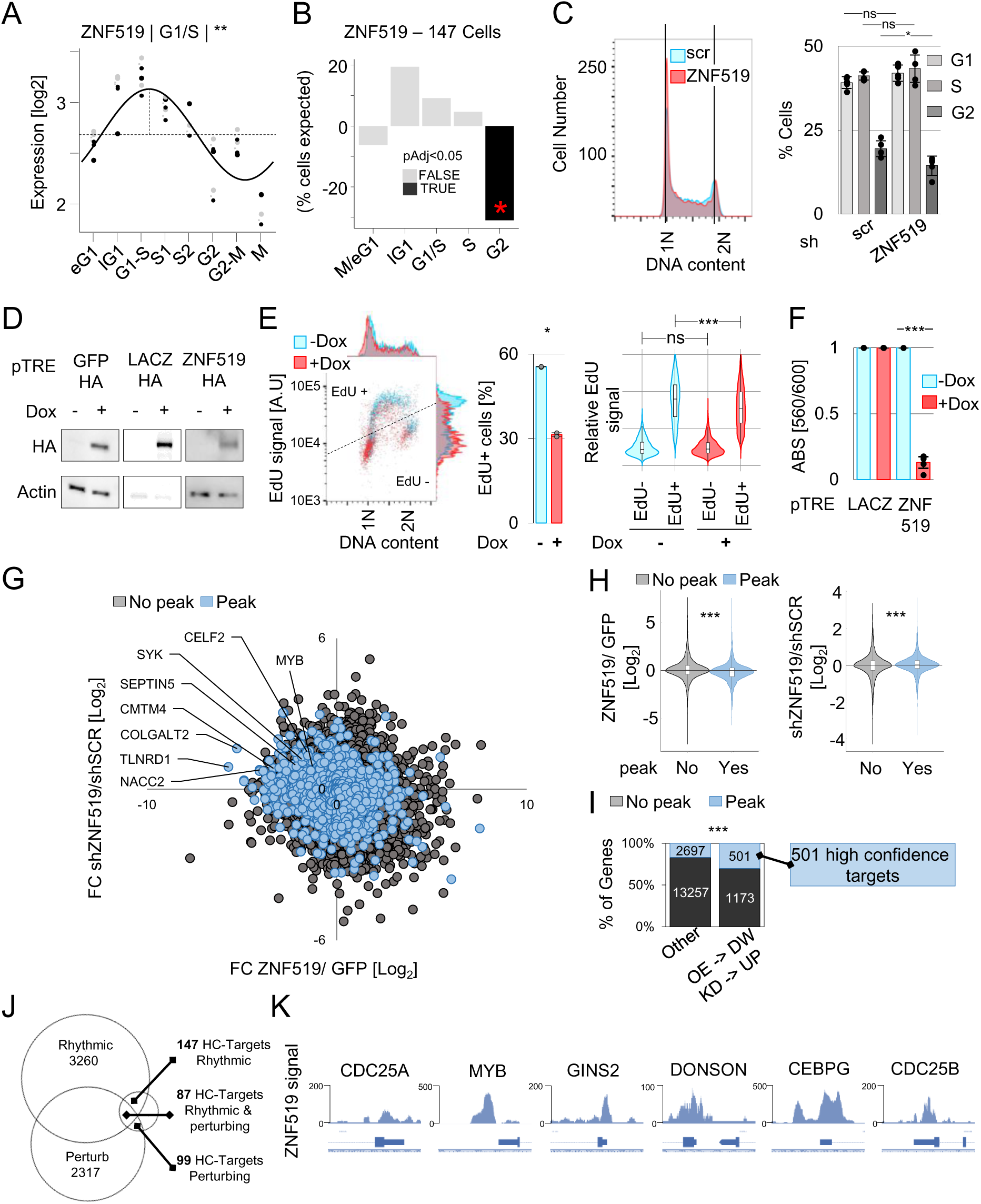
ZNF519 exerts a primate-specific layer of cell cycle regulation. (A) Cell cycle chronogram of ZNF519 expression in K562. Rhythmicity was estimated by fitting logged norm. counts to a sinusoid (adj. p-val < 0.05, F-test) (Cornelissen, 2014), with experimental batch as a covariate. Batch-corrected expression values are shown as pairs of grey (raw) and black (corrected) dots connected by a dotted curved segment. Horizontal dotted line: baseline expression. Vertical dotted line: acrophase. A rounded phase of peak expression is indicated on top. “**” indicates an adj. p-value < 0.01. (B) Cell cycle imbalances caused by ZNF519 depletion via CRISPRi (Replogle *et al.*, 2022). Cell cycle phases were estimated using *VeloCycle*. Data are presented as percentages of cells gained/lost relative to the cell cycle distribution of unperturbed K562 (Fig. S1A). Black bars highlight statistically significant imbalances (p-value < 0.05). (C) Left: Cell cycle distribution of shZNF519-versus shSCR-transduced K562 9 days post-transduction as assessed by FACS. DNA content was measured by DAPI signal. G1 and G2 populations are annotated as 1N and 2N, respectively. Right: Dean Jett model quantification of cell proportions in G1, S and G2. Bars represent mean values, error bars denote standard deviations, and dots indicate six replicates. Statistics: Two-tailed paired t-test; "ns" indicates p > 0.05 and "*" indicates p < 0.05. (D) Western blot analysis of K562 transduced with a dox-inducible HA-tagged ZNF519 or LACZ overexpression system 4 days post-induction with doxycycline. Protein extracts were probed with anti-HA antibodies, actin was used as a loading control. (E) Left: EdU-FACS analysis of K562 overexpressing ZNF519 (+DOX) or not (-DOX) for 4 days. DNA content was assessed by DAPI, and DNA synthesis by EdU incorporation. DAPI and EdU distributions are shown on the top and right of the scatter plot, respectively. The FACS threshold separating EdU+ from EdU-cells is shown as a dashed line. Middle: proportions of EdU+ cells. Bars represent mean values, error bars denote standard deviations, and dots indicate two replicates. Statistics: two-tailed paired t-test; "ns" indicates p > 0.05 and "*" indicates p < 0.05. Right: EdU signal in EdU- and EdU+ cells upon dox-induced ZNF519 overexpression. For each cell the signal is normalized by the mean EdU signal of EdU-cells. Statistics: two-tailed unpaired t-test; "ns" indicates p > 0.05 and "***" indicates p < 0.005. (F) Proliferation of K562 overexpressing ZNF519 (+DOX) or not (-DOX) for 5 days, assessed by metabolic activity using PrestoBlue. Bars represent mean values, error bars denote standard deviations, and dots indicate three replicates. Statistics: two-tailed paired t-test; "ns" indicates p > 0.05 and "***" indicates p < 0.005. (G) Log2 fold change gene expression of shZNF519 versus shSCR K562 at day 7 post-transduction (y-axis), relative to the log10 fold change gene expression of ZNF519- versus GFP-overexpressing K562 at day 5 post-dox induction (x-axis). Genes are categorized based on the presence (blue) or absence (grey) of a ZNF519 ChIP-exo peak (Imbeault, Helleboid and Trono, 2017) overlapping the TSS. (H) Left: Log2 fold change gene expression between ZNF519- versus GFP-overexpressing K562 at day 4 post-dox induction, categorized by the presence (blue) or absence (grey) of a ZNF519 ChIP-exo peak (Imbeault, Helleboid and Trono, 2017) overlapping the TSS. Statistical significance is indicated above the violin plot with a t-test, where “***” indicates p < 0.001. Right: Log2 fold change gene expression of shZNF519 versus shSCR K562 at day 4 post- transduction, categorized based on the presence (blue) or absence (grey) of a ZNF519 ChIP-exo peak overlapping the TSS. Statistical significance is indicated above the violin plot with a t-test, where “***” indicates p < 0.001. (I) Proportion of genes (1) downregulated upon ZNF519 overexpression and (2) upregulated upon knockdown, categorized based on the presence (blue) or absence (grey) of a ZNF519 ChIP-exo peak overlapping the TSS. Enrichment was computed using Fisher’s Exact Test, where “***” indicates p < 0.001. 501 genes bound by ZNF519, downregulated upon its overexpression and upregulated upon its knockdown are annotated as High Confidence (HC) targets. (J) Overlap between: 1/ genes whose depletion leads to cell cycle imbalances 2/ Rhythmic genes 3/ ZNF519 HC-targets. (K) Integrated Genome Browser (IGB) screenshots of representative ZNF519 HC-targets that are themselves rhythmic and leading to cell cycle imbalance upon depletion. ZNF519 ChIP-exo signal (Imbeault, Helleboid and Trono, 2017) is represented in blue.

### ZNF274 determines *KZFP* cell cycle expression

Having established that ZNF519 rhythmicity was required for unimpeded cell cycle progression, we next assessed whether other *KZFPs* displayed rhythmic expression patterns. The bulk RNA-seq chronogram revealed 118/337 (35%) rhythmic *KZFPs* (Fig. 4A), a fraction similar to other genes. Strikingly, rhythmic *KZFPs* more frequently peaked in G1 than other TFs (Fig. 4A-B, Fig. S4A). The rare G2-c *KZFPs* were evolutionarily older (Fig. 4C) and tended to bear variant KRAB (vKRAB) domains (Fig. S4B). Overall, *KZFP* rhythmicity pointed towards a gene family-specific and cell cycle-dependent regulatory mechanism.

**Figure 4:**
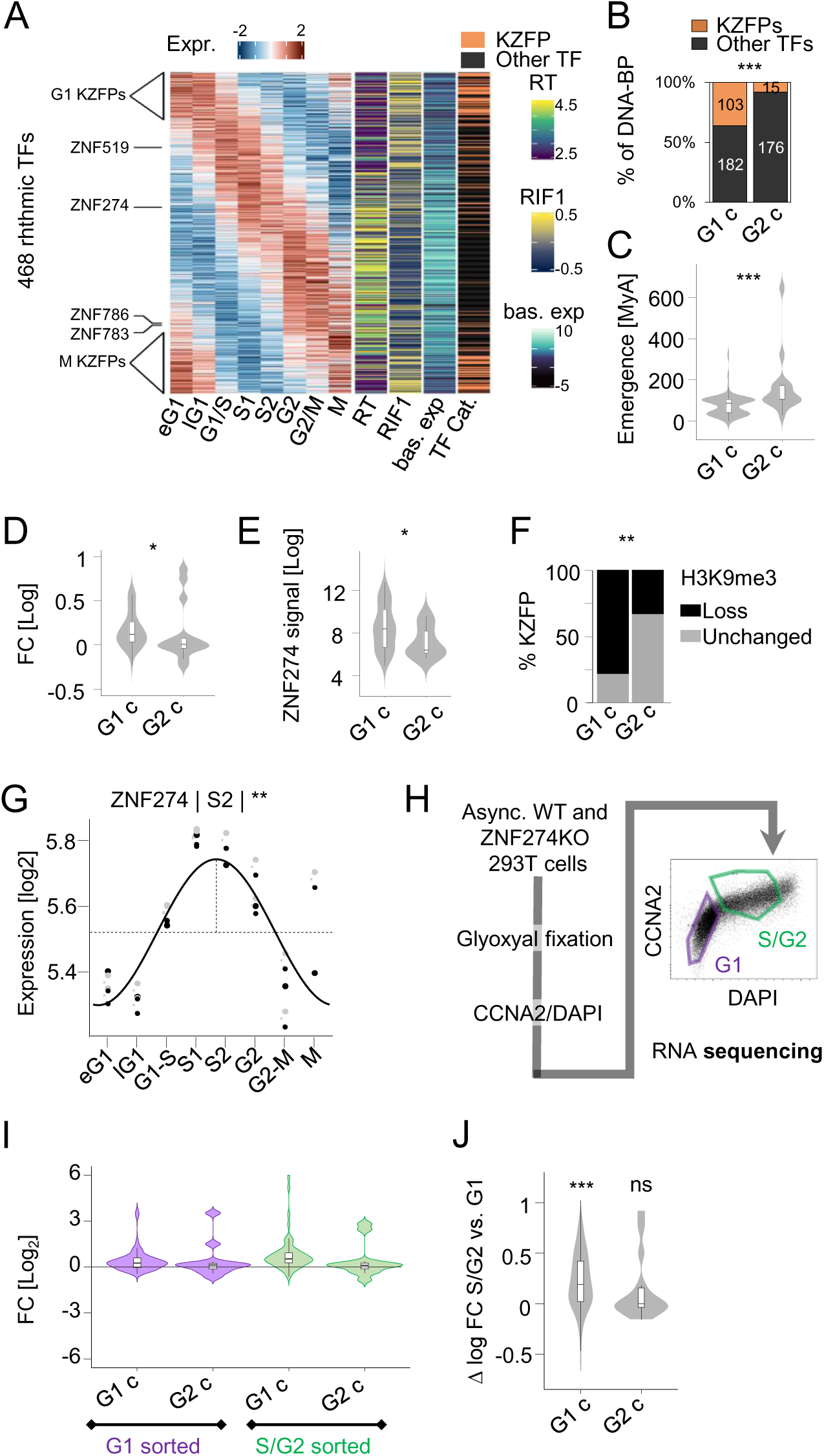
ZNF274 determines KZFP cell cycle expression. (A) Left: Z-scored expression of rhythmic TFs (adj p-value < 0.05, F-test) along the eight cell cycle bins defined in Fig. 2A. Right: 1) RT index, 2) RIF1 binding enrichment and 3) baseline expression. KZFPs are highlighted in orange. (B) Proportion of G1-c versus G2-c rhythmic KZFPs and other TFs. “***” denotes a p-value < 0.005, Fisher’s Exact Test. (C) Estimated evolutionary age of G1-c versus G2-c rhythmic KZFPs. “***” denotes a p-value < 0.005, permutation test. (D) Log2 fold change expression of G1-c versus G2-c rhythmic *KZFPs* in *ZNF274* KO versus wild-type (WT) 293T (Begnis *et al.*, 2024). “*” denotes a p-value < 0.05, Wilcoxon’s test. (E) ZNF274 ChIP-seq signal at the 3’ ZF-encoding exon of G1-c versus G2-c *KZFPs* in K562. * denotes a p-value < 0.05, Wilcoxon’s test. (F) Proportion of G1-c versus G2-c rhythmic *KZFPs* whose 3’-ZNF encoding exons overlap a region of H3K9me3 loss induced by *ZNF274* KO (Begnis *et al.*, 2024). “**” denotes a p-value < 0.01, Fisher’s Exact Test. (G) Cell cycle chronogram of *ZNF274* expression in K562. (H) Left: overview of the synchronization-free procedure for sorting G1 and S/G2 *ZNF274* KO and WT 293T cells. Right: 293T stained with CCNA2 antibody and DAPI. Sorting gates are annotated in purple (G1) and green (S/G2). (I) Log2 fold change expression of G1-c and G2-c rhythmic *KZFPs* in *ZNF274* KO versus WT 293T in FACS-sorted G1 or S/G2 cells. (J) Differential S/G2 versus G1 log2 fold change expression in *ZNF274* KO versus wild-type 293T for G1-c and G2-c rhythmic KZFPs. *** denotes a p-value < 0.005, Wilcoxon’s test with *μ* = 0.

We and others previously characterized the SCAN-containing KZFP ZNF274 as a master *KZFP* regulator ^26–29^. Specifically, ZNF274 induces H3K9me3 at *KZFPs,* thus acting as a repressor, and further directs its gene targets to nucleolus-associated domains (NADs) where silencing is reinforced ^26–29^. We therefore wondered whether ZNF274 may also control *KZFP* rhythmicity. Conspicuously, G1-c *KZFPs* underwent a collective upregulation upon *ZNF274* KO, whereas G2-c *KZFPs* remained unaffected (Fig. 4D). Moreover, G1-c *KZFPs* displayed higher ZNF274 binding (Fig. 4E) which coincided with an acute H3K9me3 loss upon *ZNF274* KO (Fig. 4F). Since *ZNF274* expression peaks in S (Fig. 4G) and given the overrepresentation of G1-c *KZFPs* amongst its targets, we hypothesized that ZNF274 may exert most of its *KZFP*-repressive activity in S to G2.

To address this, we FACS-sorted *ZNF274* KO HEK293T into G1 and S/G2 fractions before subjecting them to RNA-seq (Fig. 4H, S4B). *KZFPs* underwent a greater upregulation in S/G2 than in G1, whereas other genes exhibited no such difference (Fig. S4D, E). Moreover, in S/G2, G1-c *KZFPs* underwent a greater upregulation than G2-c *KZFPs*, whose aggregate expression remained unchanged (Fig. 4I, J). These results strongly suggest that ZNF274 – while expressed in S/G2 - represses target *KZFPs*, thus only permitting their expression in M-to-S1. Consequently, *KZFPs* escaping ZNF274-mediated repression may attain elevated G2-c expression. Altogether, these results show that ZNF274 sets the cell cycle expression of hundreds of KZFPs.

### ZNF274 sets gene-dense regions for late replication

The preferential G1-c expression of *KZFPs* coincided with a notable replication delay compared with other genes (Fig. 4A, Fig. 5A), with G2-c *KZFPs* replicating earlier than their G1-c counterparts (Fig. 4A, Fig. 5B). Unlike what we observed for other rhythmic TFs, binding of the heterochromatin-associated RT factor RIF1 did not correlate with RT at rhythmic *KZFPs* (Fig. 5C). Accordingly, *KZFP* clusters were strikingly insensitive to *RIF1* KO, thereupon retaining late replication (Fig. 5D). Together, these results hint that RT at *KZFPs* may largely be RIF1-independent.

**Figure 5:**
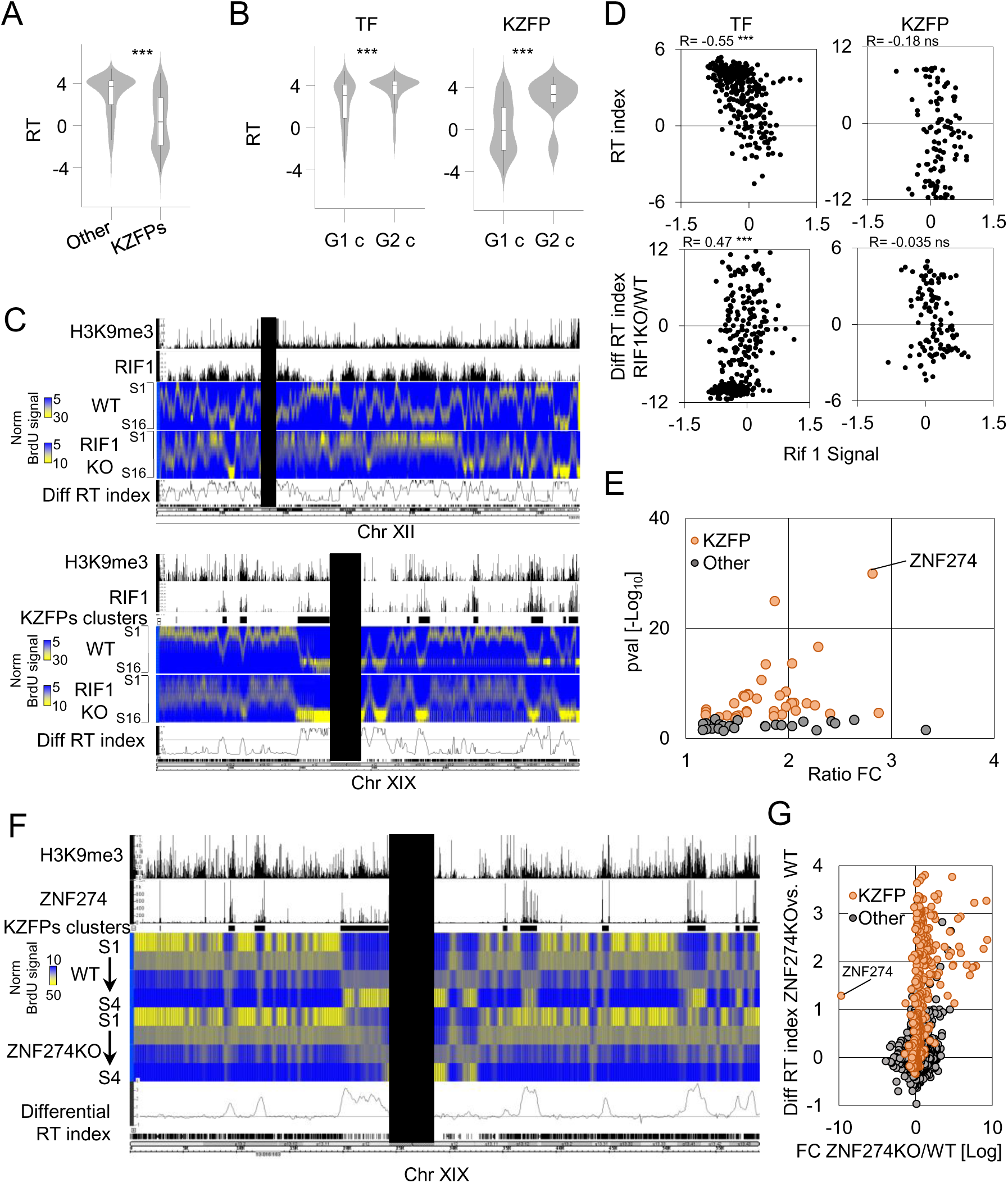
ZNF274 sets gene-dense regions for late replication. (A) RT index of *KZFPs* and other TF-encoding genes in K562. “***” denotes a p-value < 0.005, Wilcoxon’s test. (B) RT index of G1-c and G2-c *KZFPs* and other TFs in K562. “***” denotes a p-value < 0.005, Wilcoxon test. (C) IGB screenshot showing chromosomes 12 (top) and 19 (bottom). H3K9me3 ChIP-seq and RIF1 binding are depicted in black. BrdU-IP signal (percentage of repli-seq reads falling in each S phase fraction) is shown as a heatmap for WT and *RIF1*-KO HCT116, along with the corresponding differential RT index. *KZFP* clusters are represented as black rectangles. Centromeric regions have been masked. (D) RIF1 binding versus RT index in K562 (top) or differential RT index between *RIF1* KO and WT HCT116 (bottom) at rhythmic *KZFPs* or other rhythmic TFs. (E) DBP enrichment across late-replicating, *RIF1* KO-unaffected, H3K9me3-dependent regions (Klein *et al.*, 2021) (Fisher’s Exact Test, adj. pval < 0.05). (F) IGB screenshot showing chromosome 19. H3K9me3 and ZNF274 ChIP-seq signals in K562 are depicted in black. BrdU-IP signal is represented as a heatmap for *ZNF274* KO and wild-type 293T and the differential RT index between the two cell lines is reported as a black line. *KZFP* clusters are represented as black rectangles and centromeric regions have been masked. (G) Log2 fold change expression and differential RT index of KZFPs (orange) and other genes (grey) in *ZNF274* KO versus wild-type 293T.

ZNF274 affects both 3D genome organization and epigenetic states ^28,29^, two strong correlates of RT ^30^. Moreover, ZNF274 binding highly enriched within late-replicating regions unaffected by *RIF1* KO, many corresponding to *KZFP* clusters (Fig. 5C). Remarkably, late replicating RIF1-unaffected regions undergoing a late-to-early shift upon additional knockdown of the H3K9me3 methyltransferases *SETDB1*/*SUV39H1*/ *SUV39H2* revealed a striking enrichment for ZNF274 (Fig. 5E). Thus, we hypothesized that ZNF274 sets RT at its genomic targets – many of which are *KZFPs* – independently from RIF1.

To address this, we established genome-wide RT maps for *ZNF274* KO and WT HEK293T by repli- seq ^31,32^ (Fig. S5A). *ZNF274* ablation induced dramatic late-to-early RT shifts at ZNF274-targeted *KZFPs* (Fig. 5F-G). Other ZNF274 targets, such as *PCDH*s ^29^ and KRAB-devoid *ZFPs* followed the same trend (Fig. S5B). *ZNF274* KO-induced RT switches spanned well-defined, gene-rich regions, leaving the rest of the genome unaffected (Fig. 5F, G). These results show that ZNF274-instated H3K9me3 partakes in setting RT at specific gene-dense regions genome-wide.

## Discussion

Conserved genes are enriched for processes common to most living organisms, e.g. mRNA processing and cell cycle progression, whereas genes undergoing accelerated evolution are enriched for environment-interaction, e.g. immunity and sensory perception ^33^. Yet, substantial differences pertaining to multiple levels of cell cycle regulation have been documented across the tree of life. For instance, the cyclin-dependent kinase inhibitor CDKN1A is restricted to mammals ^3,20^, which split from birds and reptiles ∼250mya. This begs the question: what is the higher bound on lineage-specific cell cycle regulation?

Our work demonstrates that despite their rapid evolution, KZFPs, which encompass most of the mammalian-specific TFs, partake in multiple aspects of the cell cycle. While we did find that conserved TFs were overall more likely to regulate the cell cycle, we nonetheless identified ten evolutionarily recent TFs (emerged less than 250mya) whose binding was enriched at the promoters of rhythmic genes, and whose silencing compromised cell cycle progression. Of those, eight were KZFPs. Among them, we found that ZNF519, a primate-restricted KZFP expressed in G1/S, represses hundreds of cell cycle-rhythmic genes – many peaking in G1/S-to-S - via direct promoter binding. Much like E2F7/E2F8 ^34,35^, ZNF519 likely closes a transcriptional negative feedback loop triggered in late G1. This is further supported by widespread E2F1/TFDP1/TP53 binding at the *ZNF519* promoter across cell types. Remarkably, the absence of *ZNF519* orthologs anterior to the Gibbon-human divergence suggests that this regulatory dependency emerged rather rapidly, within less than 20MY. In contrast, *E2F7*/*E2F8* orthologs are found in *A. Thaliana*, whose last common ancestor with humans dates back to ∼1.6 billion years ago (bya). The emergence of ZNF519 in cell cycle control may thus have been driven by its transposable element targets ^15^, which appear to have adaptatively mimicked promoter sequences to multiply, thereby diverting ZNF519 binding towards promoters ^22^. Based on our systematic characterization of KZFPs throughout the cell cycle, we anticipate that several of them influence this biological process along similar means ^22^. For example, *ZNF695* silencing causes a profound attrition in S; its expression is rhythmic; its binding sites overlap human-specific indels at promoters ^22^ and its expression is strikingly associated to proliferation across cancer types ^36^. Put together, these results hint that ZNF695, a 30myo KZFP, partakes in cell cycle regulation.

Moreover, KZFPs exhibit a family-wide tendency for synchronous cell cycle expression. Our bulk RNA-seq chronogram revealed that TRIM28-recruiting, i.e. heterochromatin-instigating KZFPs are generally expressed in M-to-G1 and conversely repressed in S-to-G2. This observation is consistent with one or several of the following hypotheses. First, KZFP-instigated H3K9me3 may be detrimental during late S and G2 and therefore have been selected against. Second, KZFP-elicited H3K9me3 may partake in moulding the pre-replication chromatin landscape in M-to-G1, ahead of replication origin licensing. Interestingly, as licensing ends with G1 and given that euchromatin is licensed before heterochromatin, shortening G1 causes under-replication and genomic stress preferentially within the latter ^37^. We thus speculate that some of the cell cycle imbalances caused by *KZFP* silencing may stem from improper origin licensing resulting from an aberrant lG1 heterochromatic landscape. Along these lines, we have recently shown that depleting ZNF417 and ZNF587, two primate-specific KZFPs frequently upregulated in lymphoma cells, induces H3K9me3 loss and replication stress at their target sites ^38^. Third, the G1-c expression bias of most KZFPs may be a consequence of an adaptation at the level of RT, as late RT generally correlates with G1-c expression.

Interestingly, many of the rare G2-c KZFPs appear structurally unable to recruit TRIM28 ^16,25^, having likely specialized in H3K9me3 deposition-independent functions during evolution. Our systematic assessment of KZFP binding at rhythmic promoters has for instance revealed that the 105 myo vKRAB- bearing ZNF786 is enriched at the promoters of G1/S-to-S1 rhythmic genes. Establishing the molecular features of this vKRAB domain promises to shed light on the function of ZNF786 within the cell cycle.

The cell cycle expression of *KZFPs* hinted at a common regulatory mechanism. Our data establish ZNF274, a eutherian-restricted TF, as a master controller of *KZFP* rhythmicity. Mechanistically our results suggest that upon reaching peak expression in late S – a remarkably rare feat amongst TRIM28- recruiting KZFPs - ZNF274 binds the 3’ ZF-encoding exons of target *KZFPs* and KRAB-devoid *ZFPs* hence instigating H3K9me3. An S-to-G2 repression of target *KZFPs* ensues, which is later reduced as ZNF274 levels fade in M-to-G1. Accordingly, the ZF-encoding exons of G2-c *KZFPs* are generally ZNF274^low^. This suggests that ZNF274’s target specificity likely depends on ZF-encoding exon-located DNA-binding motifs, their integrity and density locally determining the affinity of ZNF274’s own zinc finger (ZF) array. G2-c KZFPs are evolutionarily older than G1-c KZFPs, consistent with a model whereby their ZF-encoding exons had more time to diverge from the canonical, ZNF274-affine ZF- encoding exons of more recently emerged *KZFPs*. Whether ZNF274-driven nucleolus tethering ^29^ also partakes in the process is not yet clear.

The impact of KZFPs on the cell cycle is not limited to gene regulation. We have found that ZNF274 controls RT at hundreds of genes - including *KZFPs*, *PCDHs* and select KRAB-devoid *ZFPs* – in a RIF1-insensitive manner. Several parallels can be drawn between ZNF274 and RIF1. Both simultaneously affect RT and the 3D genome structure: ZNF274 targets genes to the nucleolar periphery ^29^ while RIF1 enriches within nuclear lamina-associated domains ^39^. Both associate with H3K9me3 domains albeit of different intensities: RIF1 binds to H3K9me3-intermediate regions ^10^ while ZNF274 instigates high levels of the same mark at its target sites ^26,28,29^. Both paradoxically associate with early replication in specific contexts: mouse Rif1 associates with early RT in B cells ^40^ whereas *KZFP* clusters replicate early in embryonic stem cells ^41,42^ despite substantial H3K9me3 and ZNF274 binding ^28^, which may be explained by the transcriptionally hyperactive state of the nucleolus in that context ^43^.

However, ZNF274 and RIF1’s evolutionary trajectories differ widely. *RIF1* shares an ortholog with *S. Cerevisiae* and thus must have emerged at least ∼1 billion years ago (bya) whereas *ZNF274* only appeared 105mya ^15^. Moreover, their respective target regions suggest adaptations brought about in the face of distinct selected pressures. RIF1 sets vast, often gene-poor regions for late RT thereby divesting limiting replication factors towards early-replicating regions. Meanwhile, ZNF274 sets late RT at gene-dense regions, their summed genomic sizes likely being too short to compete with early-replicating regions upon *ZNF274* ablation. It is thus tempting to speculate that ZNF274, having emerged in the context of a RIF1-controlled RT, has evolved to exert a lineage-specific layer of RT insulated from RIF1.

ZNF274-targeted *KZFPs* generally replicate late, at least in somatic cells, which may be of functional relevance for the following reasons. First, the tandem-arrayed and hence “recombigenic” nature of *KZFPs* ^27^ – just like that of *PCDHs* - may generate replication stress. Late RT may thus favor the recruitment of a risk-aware DNA replication machinery, similarly to what has been reported at common fragile sites ^44^. Accordingly, ZNF274 binds *KZFPs* alongside ATRX ^28^, a protein involved in DNA damage prevention at repetitive regions. Moreover, *KZFPs* display elevated H3K36me3, which has been associated to DNA repair ^45^. Along the same lines, heterochromatin displays a higher risk of *cis*- translocations relative to euchromatin, where *trans*-translocations are instead more likely ^46^. Genes are generally associated with euchromatin, but not *KZFP* clusters at which a higher risk of *cis*- translocations may have been traded against the genomic threat of *trans*-translocations. Lastly, KZFPs may be targeted for late RT as a consequence of potentially selected expression patterns outlined above. As often, whether an adaptative RT constrains transcription or the reverse remains to be clarified.

Poly-zinc finger proteins such as KZFPs are ubiquitous features of metazoan genomes ^15,47^, their fast evolution matching the spread of transposable elements ^15,48–52^. Rapid evolution suggests adaptative fine tuning; indeed, KZFPs, along with their TE targets, are recurrently found necessary for the development and function of specialized tissues ^33,53–65^. However, whether such evolutionarily recent TFs partake in comparatively ubiquitous processes remained controversial. Our work sheds light on hitherto unsuspected regulatory and mechanistic ties between KZFPs and cell cycle progression, one of the most fundamental of biological processes. This paves the way for understanding how distinct genomes acclimate to the pervasiveness of cell division and the daunting selective pressure thereby exerted on DNA replication, while providing the regulatory plasticity required for adapting to an ever-changing environment. Whether the resulting transcriptional dependencies are only neutral by-products of a regulatory turnover enabling the evolvability of environment-interaction genes ^33^, or instead underly yet unidentified phenotypic changes in the conduct of an otherwise highly conserved process ^66^ remains to be established. The latter would likely be most relevant during organogenesis, when proliferation and cell fate commitment are closely intermingled.

## Materials and Methods

### Data visualization

The bulk chronogram of cell cycle gene expression, the perturbation-induced imbalances and the enrichment of DNA-binding proteins at promoters of cell cycle rhythmic genes are available for further exploration in Shiny-R at https://tronoapps.epfl.ch/cycleQuest/.

### Cell Culture Conditions

K562 and HEK293T cells were maintained in RPMI-1640 (Gibco) and DMEM (Gibco), respectively, supplemented with 10% fetal bovine serum (FBS) and 1% penicillin-streptomycin (Gibco) at 37°C in a 5% CO₂ humidified incubator. K562 cells were seeded at 1x10⁵ cells/mL and passaged every 2-3 days to maintain exponential growth. 293T cells, obtained from Begnis et al., 2024, were cultured to 80% confluence and split at a 1:10 ratio.

### Cell Cycle Fluorescence Activated Cell Sorting prior to RNAseq

Asynchronous K562/HEK293T cells were fixed according to protocol Channathodiyil and Houseley (2021). Briefly, cells were fixed using a glyoxal/methanol solution to preserve nuclear structure. Following fixation, cells were washed and stained for CCNA2 (Cyclin A2) using a primary antibody (PE anti-Cyclin A Antibody – Biolegend – REF: 644004) at a 1:200 dilution in blocking solution (1% BSA in PBS). for 1 hour at room temperature. Nuclear content was subsequently stained with DAPI (0.5 µg/mL) for 15 minutes in the dark. Cells were submitted to FACS according to the cell cycle using FACSAria Fusion system. Following sorting, total RNA was extracted.

### RNA Extraction and Quality Control

Human RNA samples were collected from four distinct experiments: CellCycle-K562, OE-ZNF519, CellCycle-HEK293T-ZNF274KO and KD-ZNF519. Total RNA was extracted from K562 cells using the NucleoSpin RNA Plus, Mini kit for RNA purification with DNA removal column from Macherey-Nagel (Ref: 740984.50) following the manufacturer’s instructions. RNA concentration and quality were assessed using a NanoDrop 2000 (Thermo Fisher) and Agilent Bioanalyzer 2100 (Agilent Technologies). RNA quality was assessed using a Nanodrop spectrophotometer (Thermo Fisher Scientific) to measure concentration and purity, followed by fragment size analysis using the TapeStation 4200 system (Agilent Technologies). When discrepancies between Nanodrop and TapeStation measurements were observed, RNA concentrations were confirmed using the Qubit RNA High Sensitivity assay (Thermo Fisher Scientific). For the CellCycle-K562 experiment, where the RNAs were partially degraded (RIN 5–6.4), corrections were applied to the 18S/28S peak assignments. Final RNA concentrations for library preparation were calculated as the average of Nanodrop, TapeStation, and Qubit measurements.

### RNA Library Preparation with NEBNext Ultra II Directional RNA Library Prep with Ribodepletion

For the CellCycle-K562 experiment, RNA libraries were prepared using the NEBNext Ultra II Directional RNA Library Prep Kit with ribodepletion (New England Biolabs, NEB protocol Version 4.0_4/21), starting from 51 ng of input RNA. Ribosomal RNA was depleted, and libraries were constructed following the manufacturer’s instructions. Library quantification was performed using the Qubit dsDNA High Sensitivity (HS) assay (Thermo Fisher Scientific), and fragment size distribution was confirmed with the TapeStation 4200. While most libraries passed quality control, two samples (RF053_S3_2 and RF054_M_3) displayed suboptimal profiles but were deemed acceptable for sequencing.

### RNA Library Preparation with Illumina Stranded mRNA Library Preparation

For the OE-ZNF519 and CellCycle-HEK293T-ZNF274KO and KD-ZNF519 experiments, RNA libraries were prepared using the Illumina Stranded mRNA Ligation Kit (Illumina, protocol 1000000124518 v03). The OE-ZNF519 libraries were constructed from 500 ng of RNA, while the CellCycle-HEK293T-ZNF274KO and KD-ZNF519 libraries were prepared from 200 ng of RNA. Poly(A) selection and directional library construction were performed according to the manufacturer’s protocol. The resulting libraries were quantified with the Qubit DNA HS assay and checked for fragment size distribution on the TapeStation 4200. All libraries met the quality and yield requirements for sequencing.

### RNA Sequencing

Sequencing of prepared libraries was conducted using the NovaSeq 6000 system (Illumina). For the CellCycle-K562 experiment, sequencing was performed in a paired-end 111 bp configuration (PE111). The CellCycle-HEK293T-ZNF274KO and KD-ZNF519 libraries were sequenced in a paired-end 60 bp configuration (PE60), while the OE-ZNF519 libraries were sequenced in a paired-end 100 bp configuration (PE100).

### Lentivector production

K562 cells overexpressing HA-tagged KZFPs were generated as described in <u>Imbeault et al. (2017)</u>. In short, cDNAs from the human KZFPs were codon-optimized and synthesized using the GeneArt service from Thermo Fisher Scientific (former Life Technologies). Sequences were cloned into the doxycycline inducible expression vector pTRE-3HA which yields C-terminally tagged proteins. Stable cell lines were generated using Lentivector transduction of mycoplasma free HEK293T cells as described on http://tronolab.epfl.ch.

### Knockdown (KD) Protocol in K562 Cells

K562 cells were transduced with lentiviral particles expressing shRNA targeting the gene of interest (shZNF519: TRCN0000428578Sequence: 5’-GCCTTACAACTCTAATGAATG-3’) or scrambled sequence (shSCR – Sequence: 5’-CAACAAGATGAAGAGCACCAAG-3’). Cells were seeded at a density of 1x10⁵ cells/mL and transduced. After 48 hours, cells were treated with 1 µg/mL puromycin for 3 days to select for successfully transduced cells. After selection, cells were replated at 1x10⁵ cells/mL and collected for further analyses at various time points up to 10 days post-transduction.

### Overexpression (OE) Protocol in K562 Cells

Doxycycline (Dox)-inducible overexpression was performed using a lentiviral system. K562 cells were transduced with lentiviral particles containing GFP-3XHA, LACZ-3XHA or codon optimized ZNF519- 3XHA constructs under the control of a tetracycline-inducible promoter. Cells were seeded at 1x10⁵ cells/mL and transduced. After 48 hours, cells were selected with 1 µg/mL puromycin for 3 days. Overexpression was induced by adding 1 µg/mL doxycycline to the culture medium for 5 days. Induction efficiency was validated by Western blot.

### Cell Cycle Assessment with Propidium Iodide (PI) Staining

Cell cycle distribution was determined using propidium iodide (PI) staining. K562 cells were fixed in 70% ethanol overnight at -20°C, washed with PBS, and resuspended in a staining solution containing 50 µg/mL PI and 100 µg/mL RNase A in PBS. Cells were incubated for 30 minutes at room temperature in the dark and analyzed on a Gallios flow cytometer (Beckman Coulter) for DNA content. Cell Cycle analysis was performed with flowjo and DeanJettFox model was used to calculate cell proportion in the different phases.

### EdU Incorporation and FACS Analysis

Proliferating cells were labeled with 10 µM EdU for 30 minutes. After labeling, cells were washed with 1% BSA in PBS and fixed with 2% paraformaldehyde (PFA) in PBS for 30 minutes at room temperature. Following fixation, cells were washed again with 1% BSA in PBS and permeabilized with 0.1% saponin in distilled water (H₂O) for 30 minutes at room temperature. After permeabilization, cells were washed with 1% BSA in PBS and subjected to the click reaction using a homemade solution containing 100 µM CuSO₄, 100 mM ascorbic acid, and 5 µM Alexa Fluor 488-conjugated azide (ref: A10266) in distilled water. The click reaction was performed for 30 minutes at room temperature, protected from light. After the reaction, cells were washed thoroughly with 1% BSA in PBS. DNA content was then stained by incubating the cells with 0.5 µg/mL DAPI and 100 µg/mL RNase A in 1% BSA in PBS for 30 minutes. Cells were washed once more with 1% BSA in PBS and analyzed using a Gallios flow cytometer (Beckman Coulter) to determine EdU incorporation and DNA content. EdU positive versus negative cells were separated according to arbitrary criteria. The EdU signal was calculated for every cell and normalized by the average of the EdU signal of the EdU negative cells.

### BrdU-IPseq (Repli-seq)

BrdU-labeled cells were fixed in 70% ethanol and stained with propidium iodide following standard protocols Zhao *et al.,* 2020. Cell sorting by DNA content was performed using a BD FACSAria SORP sorter, and 80,000 cells were collected for each of four S-phase fractions. The region between the G1 and G2 peaks was divided into four equal fractions, labeled S1–S4. Total genomic DNA was extracted from each fraction. BrdU-labeled DNA from each S-phase fraction (S1–S4) was immunoprecipitated using 0.5 *μ*g of anti-BrdU antibody (BD #555627) and 20 μg of anti-mouse IgG (Sigma #M7023) following adaptor ligation. The BrdU-DNA/anti-BrdU complexes were captured by incubating with 2 μL of Dynabeads Protein G (Thermo Fisher, #10003D) for 20 minutes at room temperature. After three washes with 200 μL of PBST (5 minutes each), the BrdU-labeled DNA was released by Proteinase K digestion and purified. NEBNext Ultra DNA Library Prep Kit for Illumina (NEB, cat. no. E7370) was used to do the library according to the manufacturer protocol with the following index set NEBNext Multiplex Oligos for Illumina (Dual-Index Primers Set 1; NEB, cat. no. E7600S) and sequenced on AVT system.

### Cell Proliferation Assay Using PrestoBlue

Cell proliferation was assessed using the PrestoBlue™ Cell Viability Reagent (Invitrogen). Briefly, 10 µL of PrestoBlue reagent was added directly to each well of a 96-well plate containing cells in 100 µL of medium. After a 2-hour incubation at 37°C, fluorescence was measured using a microplate reader following the manufacturer instruction.

### Western Blot Using HA Antibody

Proteins were extracted from cells using a homemade RIPA buffer (50 mM Tris-HCl, 150 mM NaCl, 1% NP-40, 0.1% SDS, 0.5% sodium deoxycholate, pH 7.4) and quantified using the BCA Protein Assay Kit (Pierce). Equal amounts of protein were separated on a 4-20% gradient pre-cast Tris-glycine gel from Thermo Fisher and transferred onto a nitrocellulose membrane using the iBlot 3 system from Thermo Fisher. The membrane was blocked with 5% non-fat milk in PBS-T (PBS containing 0.1% Tween-20) for 1 hour at room temperature, followed by overnight incubation at 4°C with Anti-HA-Peroxidase (High Affinity, 50 mU/ml, Roche, ref: 12013819001) and HRP-conjugated anti-actin antibody (1:5000 dilution, ThermoFisher, ref: MA5-15739-HRP) in 5% non-fat milk in PBS-T. After three washes with PBS-T, protein bands were visualized using ECL detection reagent (Advansta Inc, ref: K-12049-D50) and imaged using a chemiluminescent imaging system (Fusion FX from Vilber). Actin was used as a loading control to normalize protein expression levels.

### Software and Data Analysis

Data analyses and visualization were performed using R (R Foundation for Statistical Computing) and Microsoft Excel. Flow cytometry data were analyzed using FlowJo (BD Biosciences), and genome alignment and sequencing visualization were performed using Integrated Genome Browser (IGB). Analysis of transcription factor network at the TSS of ZNF519 was performed with metascape and MCODE tools.

### Bulk RNA-seq mapping and preprocessing

Reads were mapped to the human (hg19) genome using Hisat2 v2.l.0 ^67^. Counts on genes were generated using featureCounts v1.6 ^68^ and only uniquely mapped reads with MAPQ > 10 were kept. Normalization for sequencing depth was done using the TMM method ^69^ as implemented in the limma package of Bioconductor ^70^ and only genes that had 3 reads in at least one sample were kept for downstream analysis. Batch-adjusted normalized counts, only used for plotting purposes, were obtained by fitting a linear model batch as an explanatory variable for each gene, and subsequently removing the effect of the batch variable.

### Bulk RNA-seq chronogram of cell cycle expression

Batch-corrected counts for all samples were projected on the first three principal components, revealing two outlier samples (S3_2 and M_2) which were excluded from further analysis. For each protein-coding gene, uncorrected, TMM-normalized expression values (on the log scale) were used to fit the following cosinor function: 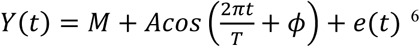 , with *Y*(*t*) the expression at time *t ε* {0, 1, … 7} corresponding to FACS-sorted cell cycle bins; *M* the Midline Statistic Of Rhythm (MESOR) i.e. baseline expression; *A* the amplitude; *T* = 8 the period; and *ϕ* the acrophase in radians capturing the time differential between the moment of maximal expression and the start of each cell cycle (*t* = 0). The equation can be rewritten as *Y*(*t*) = *M* + βx + γz + 𝛿 + 𝑒(*t*), where 𝛽 = *A*𝑐𝑜𝑠(*ϕ*); 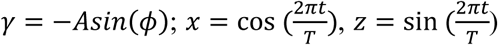 and 𝛿 encodes the sequencing batch. The model was fit using least squares and rhythmicity was determined using the F-test, comparing the full model to the reduced model *Y*(*t*) = *M* + 𝛿 + 𝑒(*t*), adjusting for multiple testing using the Benjamini-Hochberg procedure (𝑝_𝑎𝑑𝑗_ < 0.05). In the text and figures, acrophases were reported after being re-scaled as 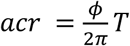. We modified the row_cosinor function from the matrixTests R package (<u>10.32614/CRAN.package.matrixTests</u>) to accept batch covariates and perform model comparisons.

### H3K9me3 promoter occupancy

Promoters were defined as previously described ^65^, i.e. as clusters of transcription start sites (derived from ENSEMBL release 93 using Biomart) spaced by less than 1 kb and extended by 500 bp at their 5′ and 3′ ends. A single promoter was assigned to each gene by sorting promoters according to decreasing H3K4me3 and H3K27ac ChIP-seq signal (bigwig, in that order) as measured in K562 by ENCODE ^11,12^, keeping the top-ranked promoter per gene. K562 ENCODE H3K9me3 ChIP-seq signal over promoters was summarized and median-scaled over 300 bins of size 10bp using deepTools ^71^ computeMatrix, extending promoters by 1.5kb on each side, thus creating a matrix with rows as genes and promoter positions as columns. The matrix was subset to promoters of rhythmic genes, reordered by acrophase and symmetrically padded on each side by a length of 7 before being median-filtering (kernel size = 4). For figs. 1F and S1H, the H3K9me3 signal was then expressed as a fraction of each promoter covered by signal exceeding the average smoothed signal across all promoters between positions 85 and 215 of the smoothed matrix. For fig. 1B, a similar fraction was computed, instead taking the median smoothed signal across all promoters as a cutoff and following an additional gaussian filtering step (kernel size = 7).

### Estimation of cell cycle phases in scRNA-seq data

The manifold-learning model from the *VeloCycle* framework ^8^ was used to infer two properties from sc/snRNA-seq datasets: (1) a continuous cell cycle phase position between 0 and 2π for single cells and (2) coefficients of a Fourier series equation describing gene-specific periodic oscillations along the cell cycle. For snRNA-seq datasets from the HL60 (702 cells), K562 (703 cells), SUDHL4 (557 cells), and THP1 (657 cells) cell lines, manifold-learning was performed jointly on all cells using the following filtering [large gene set of 376 genes, gene-wise mean unspliced level > 0.1, gene-wise mean spliced level > 0.01] and training parameters [5,000 training iterations] as recommended in the *VeloCycle* manuscript.

For analysis on the genome-wide perturb-seq dataset of K562 cells ^23^, a transfer learning approach was applied. Condition-independent estimation of the periodic Fourier series components would be especially challenging on Perturb-seq knockdown conditions containing either (1) very few cells or (2) cells belonging to just one phase of the cell cycle. To infer accurate cell cycle phases for these cells, we first performed manifold-learning for 5,000 training steps to estimate the gene harmonic coefficients (ν0, ν1sin, ν1cos) on a larger set of non-targeting control (NT) K562 cells (75,328 cells), which are more evenly distributed throughout the various phases of the cell cycle. Next, we ran manifold-learning again for 5,000 training steps, but on the entire perturb-seq dataset of 1,971,608 cells and 4,127 gene knockdown conditions (with at least 75 cells per condition). This time, we conditioned *VeloCycle* on the gene harmonic coefficients learned in the first step. This allowed cells belonging to each stratified gene knockdown condition to be assigned to a position on the cell cycle manifold, while restricting those assignments such that they were based on gene expression patterns earned on a larger and more informative dataset (allowing for batch effect expression differences).

### Correlation between bulk RNA-seq acrophases and the scRNA-seq cell phases

We circularized spearman’s rank correlation as follows: 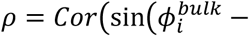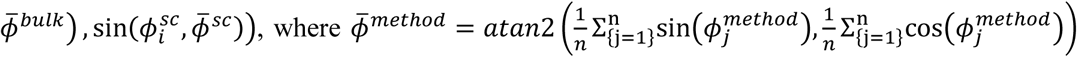 and 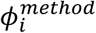 is the estimated cell cycle phase of gene 𝑖 given a method (bulk RNA seq followed by cosinor analysis, scRNA-seq followed by *VeloCycle*) in radians.

### Gene set overrepresentation analysis

Gene set overrepresentation analyses were performed using the enrichGO function from R package clusterProfiler ^72^, using the biological processes ontology, a Benjamini-Hochberg adjusted p-value cutoff on the hypergeometric test of 0.05, a minimum gene set size of 10 and a maximum gene set size of either 500 (perturbation-induced imbalances) or 100 (all other analyses). Gene universes were systematically set according to the specific dataset at hand, e.g. all genes for which rhythmicity was tested for the bulk RNA-seq chronogram, all DNA-binding proteins with available genome-wide binding data for transcriptional regulators, etc.

### Enrichment of DNA binding proteins in phase-specific rhythmic genes

DNA-binding protein ChIP-seq peaks in K562 ^11,12^ and C2H2-ZF ChIP-seq/ChIP-exo peaks in HEK293T ^13–16,73^ were overlapped with gene promoters defined as clustered transcription start sites less than 1kB away from each other and extended by 500Bp on both ends ^65^ using bedtools intersect ^74^ with default parameters. We computed enrichments of rhythmic genes assigned to each of the eight phases [eG1, lG1, G1/S, S1, S2, G2, G2/M, M] amongst those whose promoters overlapped peaks of the DNA-binding protein at hand, taking all genes on which rhythmicity test was performed in the bulk RNA-seq chronogram as a universe. Enrichment was assessed using Fisher’s Exact Test and p-values were adjusted using the Benjamini Hochberg procedure. For each DNA binding protein and each phase, specificity was defined as the median rank of adj. p-values across phases (the smallest adj. p-value receives rank 1) divided by the adj. p-value rank in the phase at hand. For ranking, ties were distributed by averaging. DNA-binding proteins with particularly statistically significant enrichments in a single phase relative to the others thus received high specificity scores.

### Gene ages

The time of emergence of genes was compiled from three sources. For *KZFPs*, we leveraged existing ab initio *KZFP* gene annotations ^73^ performed on the genomes of 69 species spanning primates, mammals, reptiles, birds and fishes. We defined the age of each *KZFP* as that corresponding to the root of the largest phylogenetic subtree whose leaves (species) contained least one paralog *KZFP* with 60% amino acid identity ^15^ at the 4-residue zinc fingerprint. For all other genes, the human-centric, vertebrate-restricted resource genTree ^20^ was used as a first pass. Genes assigned to the oldest branch of genTree were subsequently aged using GenOrigin ^19^ if the GenOrigin age was greater than the age assigned to the oldest genTree branch. Otherwise, the genTree age was used.

### Perturbation-induced imbalances

*VeloCycle*-estimated cell cycle phases *ε*[0; 2𝜋] were binned into five intervals, choosing breaks to capture the main cell cycle phases (as in G1, G1/S, etc.) and naming bins according to the gene harmonics of cell phase marker genes.

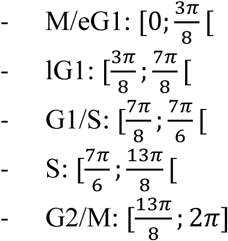

The number of breaks was chosen as a tradeoff between statistical power (see the imbalance detection test described below) and resolution, given that most perturbed conditions in the K562 perturb-seq ^23^ encompass ∼100-300 cells, leaving ∼20-60 cells in each bin. The proportions of cells from the non-targeting condition falling within each bin were used as event probabilities of a reference multinomial distribution. For each perturbed condition, the reference multinomial was sampled 1e5 times, with the number of trials set to the number of cells in perturbed condition. For each of the five bins, two proportions were computed:

- 𝑝_𝑔𝑟𝑒𝑎*t*𝑒𝑟_ the fraction of samples whereby *more* cells were observed in the bin of the perturbed condition than in the realizations of the reference multinomial

- 𝑝_𝑠𝑚𝑎𝑙𝑙𝑒𝑟_ the fraction of samples whereby *less* cells were observed in the bin of the perturbed condition than in the realizations of the reference multinomial

For any given bin, 𝑝_𝑔𝑟𝑒𝑎*t*𝑒𝑟_ ≤ 0.05 was interpreted as a cell accumulation whereas 𝑝_𝑠𝑚𝑎𝑙𝑙𝑒𝑟_ ≤ 0.05 was interpreted as a cell attrition. The matrix with vectors of relative cell proportions in each bin (rows) across conditions (columns) was projected on a 2D UMAP ^75^.

### Differential expression analysis

Differential expression analysis was performed using Voom ^76^ as implemented in the R package limma of Bioconductor v3.13. A moderated t-test (as implemented in limma) was used to test for significance. P-values were adjusted for multiple testing using the Benjamini–Hochberg procedure.

### Correlating KZFP expression to proliferation across cancer types

A proliferation signature containing a manually-curated reference set of 167 cell cycle marker genes chosen to cover all cell cycle phases was scored on each tumor sample in TCGA. To do so, counts per million of signature genes were logged, z-score-transformed per cancer subtype and averaged to a single proliferation score per tumor sample. For each *KZFP,* Spearman rank correlations were computed between *KZFP* expression values and proliferation scores in each TCGA cancer type. P-values were adjusted for multiple testing using the Benjamini-Hochberg procedure. A correlation with proliferation was called significant when 𝑎𝑑𝑗. 𝑝 < 0.05.

### Canonical KRAB domain score

To predict the canonical KRAB domain silencing potential of KZFPs, we generated a Position-Specific Scoring Matrix (PSSM) based on data from the deep mutational scan of the KRAB domain ^25^. This matrix was constructed by using HT-recruit measurements (average d5) for all possible single amino acid mutations of the ZNF10 KRAB domain. The mutation values were normalized to the wild-type construct to calculate a loss-of-silencing score. Human KRAB domains were aligned to the ZNF10 consensus sequence using MAFFT, and the aligned positions were scored for each KZFP. Computed for each KZFP, the predicted minimal score represents the mutation predicted to cause the greatest loss of silencing. For KZFPs encoding multiple KRAB domains, either as a result of alternative splicing or due to the gene structure, scores were averaged.

### Repli-seq

The raw sequencing reads were aligned to the human genome assembly hg19 using Bowtie2 ^77^ in sensitive-local mode. Subsequently, the resulting BAM files containing the aligned reads were converted into bigWig files using bamCoverage from the deepTools suite with option binSize=50000.

### RT index

The Repli-seq signal was processed as previously described ^10^. Briefly, a matrix containing the Repli-seq signal with rows as 50kb consecutive genomic windows, columns as samples and entries as the read count in the window was defined for each chromosome, left-side and right-side padded by a length of 1 and subsequently gaussian-smoothed (kernel size = 1). The resulting matrices were normalized row-wise such that each entry contains the proportion of DNA being replicated at each time point, expressed as a percentage.

The RT index was computed as previously described ^10^ as the ratio of dot products 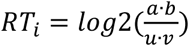 with 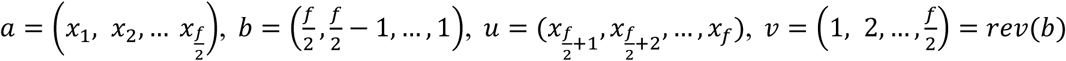 and 𝑥 = (𝑥_1_, 𝑥_2_, … , 𝑥_𝑓_) containing the percentages of DNA replicated in each of the 𝑓 FACS fractions for the given genomic region. 𝑅*T*_𝑖_ is only well-defined given an even 𝑓, and summarizes the replication timing as a single unitless value whose maximum corresponds to the earliest measured replication timing. A region with an equal proportion of replication undergoing in all 𝑓 fractions thus gets a 𝑅*T*_𝑖_ = 0.

### Identifying putative RIF1-independent, H3K9me3-dependent replication timing factors

Late-unchanged (LU) regions upon *RIF1* KO were identified as previously described ^10^. Specifically,

𝑅*T*_𝑖_ of HCT WT cells were subtracted from those of *RIF1* KO HCT cells, yielding one 𝑅*T*_𝑑𝑖𝑓𝑓_ value for each 50kb genomic window. 𝑅*T*_𝑑𝑖𝑓𝑓_ were clustered by fitting a 3-component gaussian mixture model and assigning each 𝑅*T*_𝑑𝑖𝑓𝑓_ (and thereby genomic window) to the most likely component. Genomic windows assigned to the gaussian component whose estimated mean was the closest to zero, and with 𝑅*T*𝑑𝑖𝑓𝑓 < 0 in HCT WT cells were defined as LU. LU regions were further classified according to the 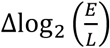 values obtained by subtracting the 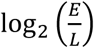 of the two-fraction repli-seq performed in *RIF1* KO sh-scramble HCT cells from that of the two-fraction repli-seq performed in *RIF1* KO *SUV39H1*/*SUV39H2*/*SETDB1* KD HCT cells. Those with 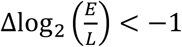 were further defined as late-to-later (LtLr), those with 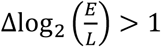 as late-to-earlier (LtEr) and those with 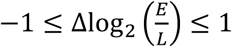 as unchanged. We defined the following genomic window categories:

- LU_K9LtEr: Late-unchanged upon *RIF1* KO, switching to an earlier RT upon additional *SUV39H1*/*SUV39H2*/*SETDB1* KD, i.e. *RIF1*-insensitive, H3K9me3-dependent.

- LU_K9LtLr: Late-unchanged upon *RIF1* KO, switching to an even later RT upon additional *SUV39H1*/*SUV39H2*/*SETDB1* KD, i.e. *RIF1*-insensitive, H3K9me3-inconsistent.

We computed statistical enrichments for LU_K9LtEr and LU_K9LtLr regions within peaks for each DNA-binding protein with available genome-wide binding data in K562 in ENCODE^11,12^ and each C2H2-ZF assayed in HEK293T cells ^13–16,73^, taking all genomic regions as a universe and using Fisher’s Exact Test and adjusting for multiple testing using the Benjamini-Hochberg procedure. Putative RIF1- independent, H3K9me3-dependent replication timing factors were defined as those whose peaks were enriched at LU_K9LtEr but not LU_K9LtLr regions.

## Data availability

Data and materials availability: All data needed to evaluate the conclusions in the paper are present in the paper and/or the Supplementary Materials. Source data have been deposited in NCBI’s Gene Expression Omnibus and are accessible through the GEO series accession number GSE280595, reviewer access: sfsdiiicvdknxut.

A website reporting resources produced with this work has been created and is publicly accessible: https://tronoapps.epfl.ch/cycleQuest/

*VeloCycle*-estimated ^8^ cell cycle phases for the K562 perturb-seq ^23^ have been deposited on ZENODO at DOI: 10.5281/zenodo.14024411

## Code availability

The code and data required to generate the figures is available at https://github.com/PulverCyril/cell_cycle

## Tables

**TableS1_GOBP:** Table with all the Gene Ontology analyses performed in the study.

**TableS2_CCExpression :** Table with the analysis of gene expression during the cell cycle.

**TableS3_GeneMetadata:** Table with the gene metadata extracted from various published studies used for the analysis.

**TableS4_PerturbSeq:** Table with the Perturb-seq RNA sequencing data.

**TableS5_OverlapPerturbXRhythmicity:** Table overlapping the analyses from Perturb-seq and the cell cycle transcriptome.

**TableS6_DNABPEnrichment:** Table with the results of the DNABP enrichment analysis over the genes based on their rhythmicity during the cell cycle.

**TableS7_OverlapPerturbXRhythmicityXDNABPEnrichment:** Table overlapping the analyses from Perturb-seq, the cell cycle transcriptome, and the DNABP enrichment at the promoters of rhythmic genes.

**TableS8_RNA-seqZNF519OE:** Table reporting the results of differential expression (DE) for genes upon the overexpression (OE) of ZNF519 in K562 cells.

**TableS9_RNA-seqZNF519KD:** Table reporting the results of differential expression (DE) for genes upon the knockdown (KD) of ZNF519 in K562 cells.

**TableS10_RNA-seqZNF274KOCellCycleSorted:** Table reporting the normalized counts for genes upon the knockout (KO) of ZNF274 in HEK293T cells and following FACS sorting into two populations according to the cell cycle.

**TableS11_Reanalysis_RepliseqRIF1KO:** Table reporting the re-analysis of the Repli-seq data from RIF1-KO (Klein et al., 2021).

## Attributions

CP, RF and DT designed the research plan. RF designed and conducted all wet lab experiments, with technical help from CR and SO. CP designed and oversaw the statistical analyses. ARL ran *VeloCycle* on the K562 perturb-seq, with input by RF, AD and EP. MB produced the *ZNF274* KO HEK293T cell line. OR provided KZFP structural metadata. JC-F analyzed the four-cell line scRNA-seq. JD and JC- F analyzed TCGA, with input by FM and RF. EP built the R Shiny App from code and data provided by CP. CP and AC performed phylogenetic analyses. JD, EP and JC-F preprocessed the raw NGS data. CP and RF analyzed the data and designed the figures. CP wrote the manuscript, with substantial contributions from RF and DT and suggestions by all authors.

## Supporting information

IndexTable

## Acknowledgements

This work was supported by grants from the European Research Council (KRABnKAP, No. 268721; Transpos-X, No. 694658), the Swiss National Science Foundation (310030_152879 and 310030B_173337), and the Aclon Foundation to D.T. We thank Philippe Pasero, Benjamin Pardo, Yea-Lih Lin, Cédric Feschotte, and Jérôme Poli for corrections and fruitful discussions. We also thank our colleagues from the Tronolab for their inputs throughout the project, and Séverine Reynard for administrative assistance.

---

**Figure Supp 1:**
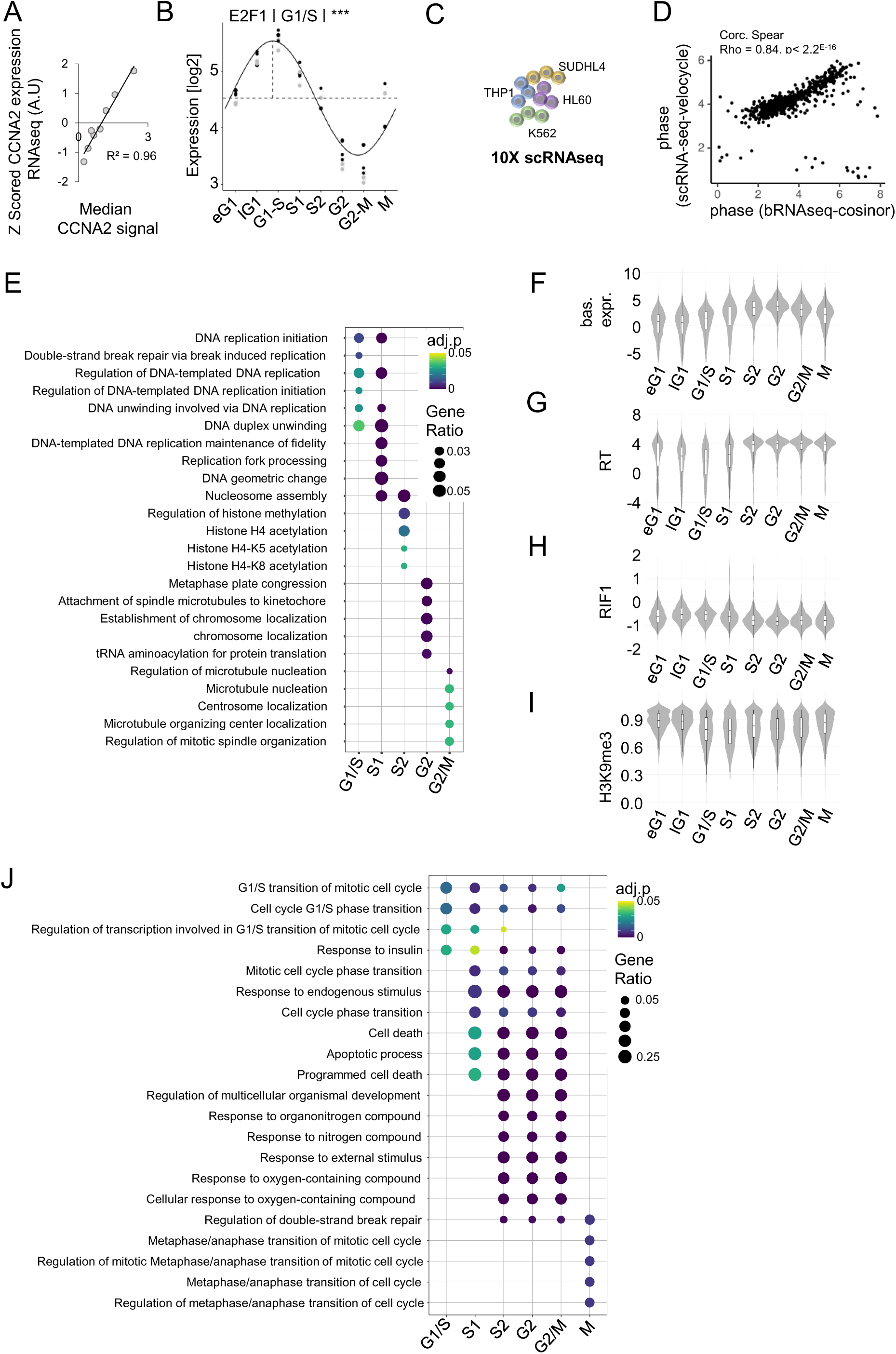
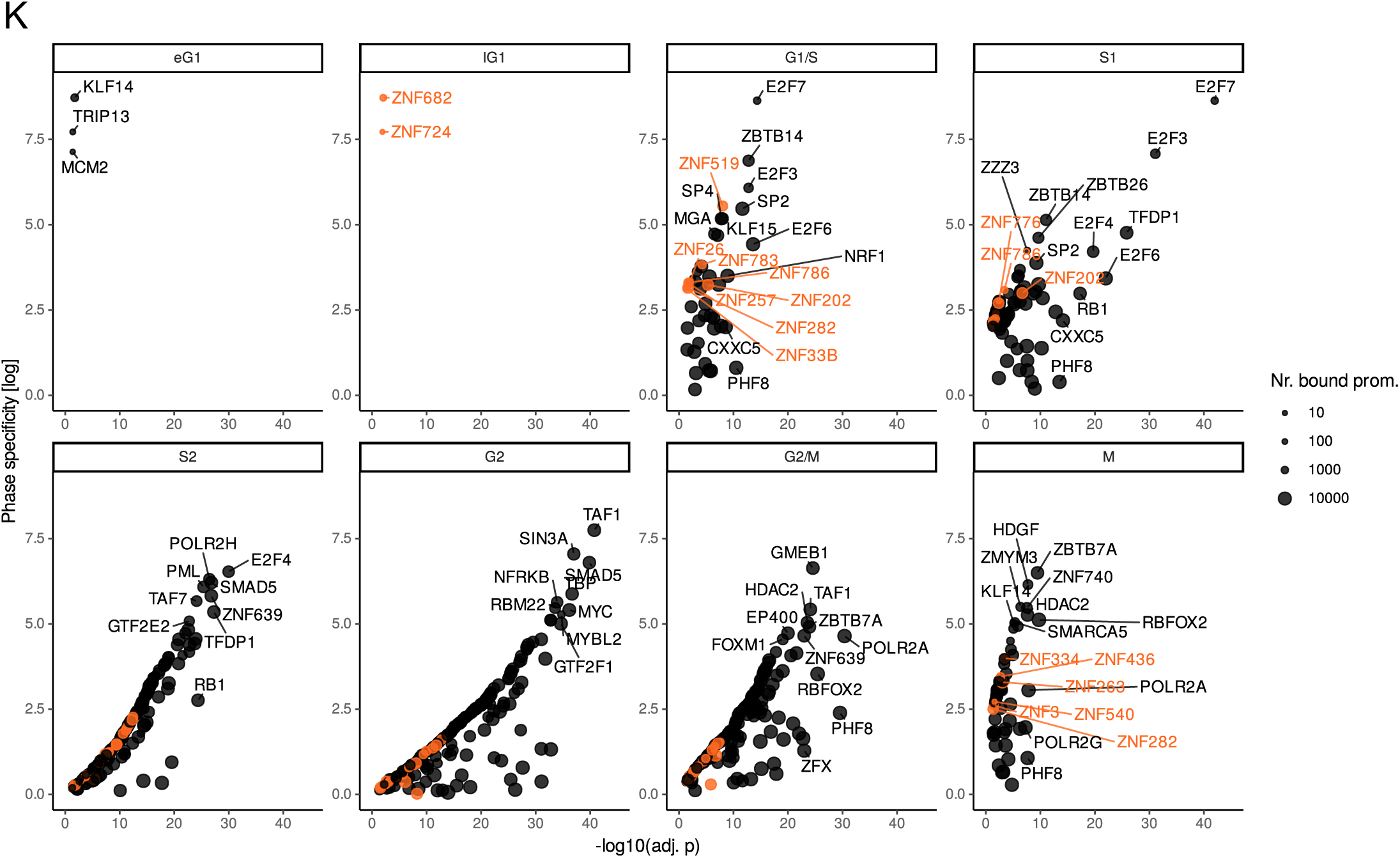
related to Fig. 1. (A) Correlation between CCNA2 expression (RNA-seq) and CCNA2 signal (FACS) across the eight gates defined Fig. 1A. (B) E2F1 expression in K562. Rhythmicity was estimated by fitting E2F1 logged norm counts to a sinusoid (Cornelissen, 2014), with experimental batch as a covariate. Batch-corrected expression values are shown as pairs of grey (raw) and black (corrected) dots connected by a dotted curved segment. Horizontal dotted line: baseline expression. Vertical dotted line: acrophase. A rounded phase of peak expression is indicated on top, with the adj. p-value of the corresponding rhythmicity F-test. “***” indicates adj. p < 0.005. (C) scRNA-seq performed on an untreated mix of four cell lines (Coudray *et al.*, BioRvx). Cell cycle phases were estimated using *VeloCycle* (Lederer *et al.*, 2024). (D) Circularized Spearman’s rank correlation between phases of peak expression estimated from the bulk RNA-seq chronogram of cell cycle gene expression and the scRNA-seq in (C) using *VeloCycle*. (E) Enrichment of rhythmic genes across Gene Ontology (GO) Biological Process (BP) terms. Dot areas represent the proportion of rhythmic genes peaking in each phase found across the enriched GO terms, and colour scale the Benjamini-Hochberg adjusted p-values (hypergeometric test). (F) Baseline expression, RT index (G), RIF1 binding (H) and H3K9me3 promoter coverage (I) of rhythmic genes across phases. (J) Enrichment of phase-enriched DBPs across GOBP terms. (K) Phase-specific enrichment of DBPs across rhythmic promoters. Dot areas represent the number of genes whose promoters are bound by the DBP. KZFPs are highlighted in orange.

**Figure Supp 2:**
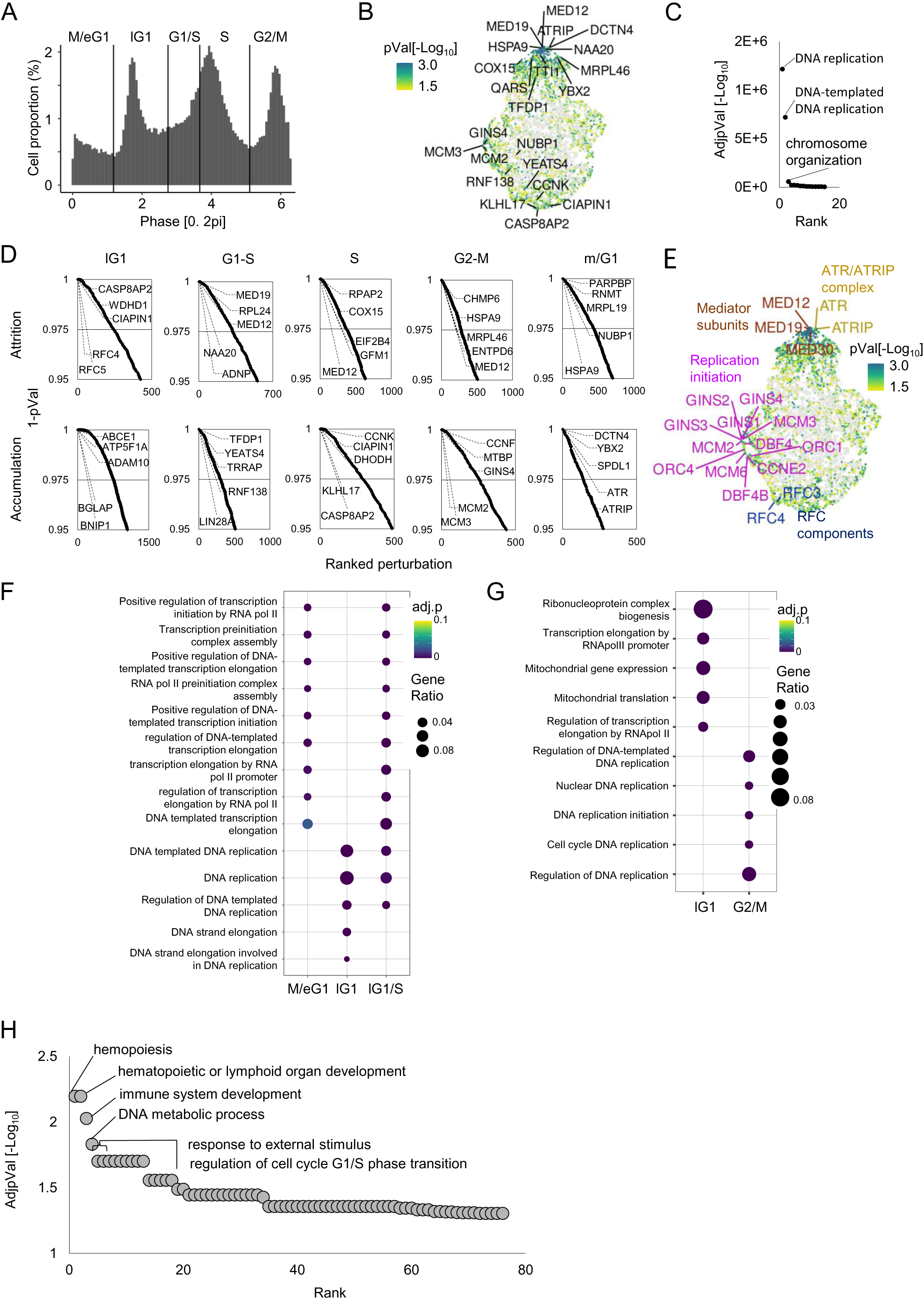
Relates to Fig. 2. (A) Cell cycle phase distribution of K562 transduced with non-targeting guide RNAs (Replogle *et al.*, 2022) estimated using *VeloCycle* (Lederer *et al.*, 2024). Phases were further split into five bins (M/eG1, lG1, G1/S, S, G2/M). (B) UMAP projection of cell cycle imbalances calculated from a K562 perturb-seq (Replogle *et al.*, 2022) with *VeloCycle* (Lederer *et al.*, 2024). Each dot corresponds to one perturbation and is colored by the statistical significance of a test based on a multinomial with parameters derived from a binned, unperturbed cell cycle phase distribution (Fig. S1A). Non-significant imbalances (p>0.05) are shown in grey. The three genes leading to the most severe imbalances (either accumulation or attrition, smallest p-values) in each phase are highlighted. (C) Enrichment of imbalance-inducing (p < 0.05) perturbation targets across GOBP terms, ranked by statistical significance (Benjamini-Hochberg adj. p-values, hypergeometric test) (D) Perturbations inducing cell attritions (top) or accumulations (bottom) (p < 0.05) in the bins shown in (A). (E) Same UMAP as in (Fig. S2B), highlighting perturbations inducing imbalances and targeting components of the same functional complexes or pathways. (F) Enrichment of perturbations leading to attritions or accumulations (G) across GOBP terms. (H) Enrichment of DBPs matching the following three criteria: (1) causing cell cycle imbalance when depleted, (2) enriched at the promoters of rhythmic genes, and (3) encoded by significantly rhythmic genes across GOBP terms (hypergeometric test).

**Figure Supp 3:**
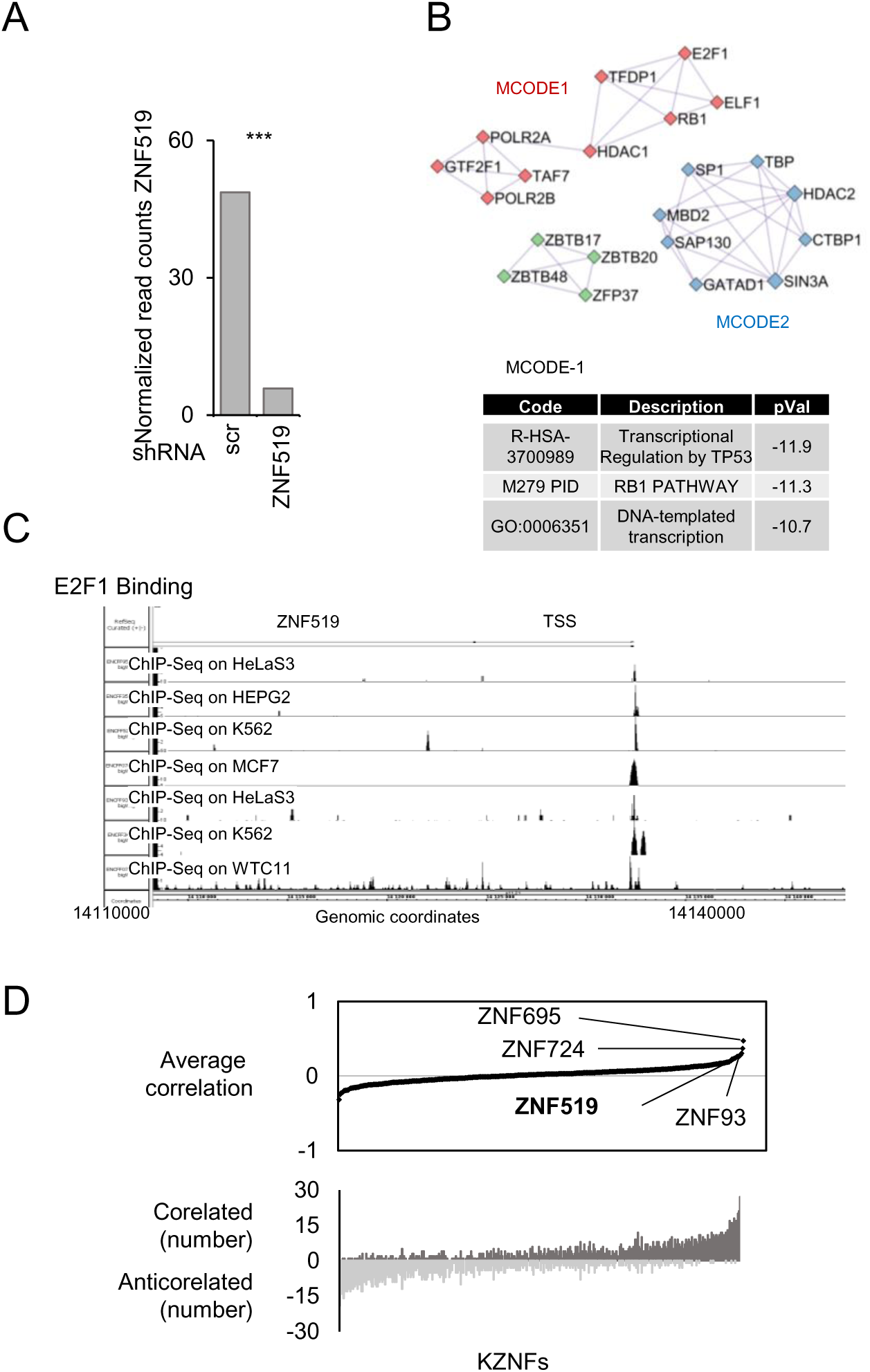
Relates to Fig. 3. (A) ZNF519 expression (RNA-seq) in shZNF519 versus shSCR K562 at day 4 post-transduction. Bars represent mean normalized RNA counts. n = 3, *** denotes a p-value < 0.005, moderated t-test. (B) Protein-Protein Interaction Network built using DBPs bound to the promoter region of *ZNF519* (GeneHancer Identifier: GH18J014131, TSS distance = +0.1 Kb) and analyzed with the mature complex identification algorithm, which allows for the identification and annotation of functional protein complexes. The table reports GO terms associated with the subnetwork MCODE1. (C) IGB screenshot showing the *ZNF519* TSS. E2F1 binding measurements conducted in various cell lines are depicted in black (The ENCODE Project Consortium, 2012). (D) Top: Median Spearman’s rank correlation of KZFP expression with 168 cell cycle and proliferation markers across 33 TCGA cancer subtypes. Bottom: Number of cancer subtypes in which KZFP expression is correlated (black) or anticorrelated (grey) with the proliferation signature (pVal <0.05).

**Figure Supp 4:**
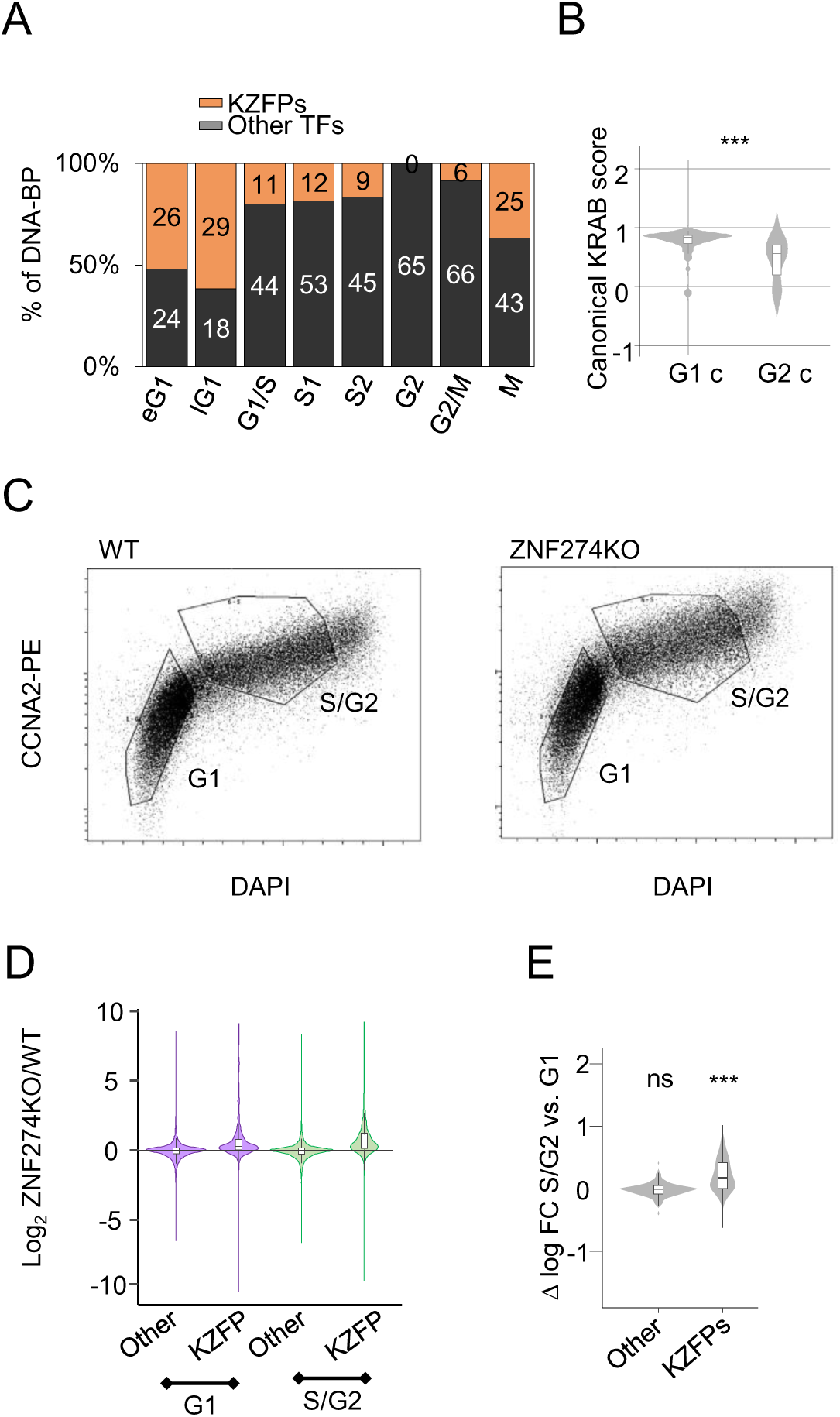
Relates to Fig. 4. (A) Proportion of rhythmic KZFPs and other TFs peaking in the different cell cycle phases. (B) Prediction of the silencing strength (1 means optimal silencing) exerted by KRAB domains of G1-c and G2-c rhythmic KZFPs. (C) *ZNF274* KO and wild-type 293T stained with anti-CCNA2 antibody and DAPI. G1 and S/G2 sorting gates are depicted in black. (D) Log2 fold change expression of rhythmic KZFPs and other genes in *ZNF274* KO versus wild-type 293T in FACS-sorted G1 or S/G2 cells. (E) Differential S/G2 versus G1 log2 fold change expression of rhythmic KZFPs and other genes in *ZNF274* KO versus wild-type 293T . “***” denotes a p-value < 0.005, Wilcoxon’s test with *μ* = 0.

**Figure Supp 5:**
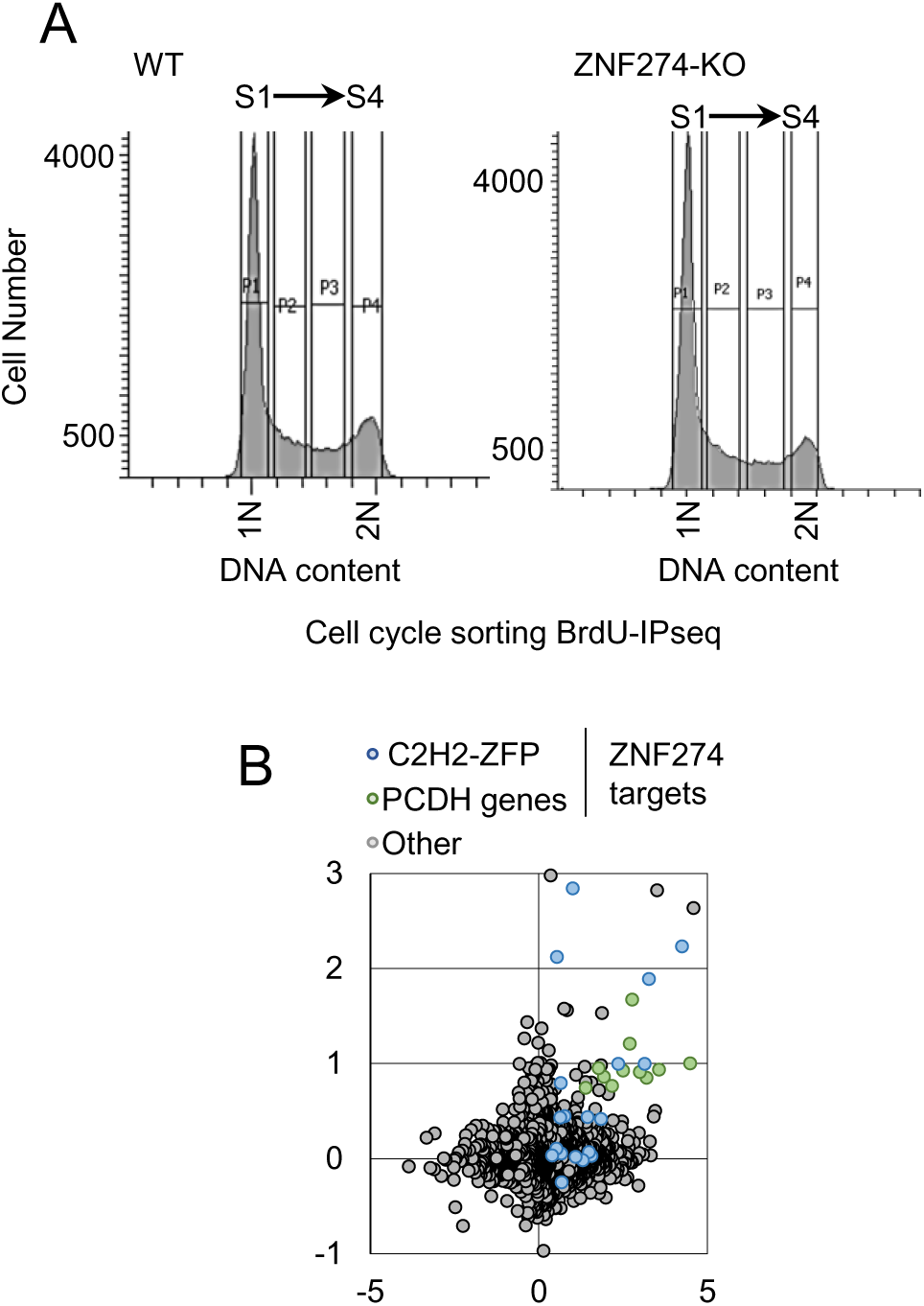
Relates to Fig. 5. (A) Cell cycle distribution of *ZNF274* KO and WT 293T stained with propidium iodide (PI). Sorting gates are depicting and annotated in black. (B) Log2 fold change expression and differential RT index of C2H2 ZFP (blue) and PCDH genes (green) targeted by ZNF274, and other genes (grey) in *ZNF274* KO versus WT 293T.

